# Activatable zymography probes enable in situ localization of protease dysregulation in cancer

**DOI:** 10.1101/2020.07.29.227413

**Authors:** Ava P. Soleimany, Jesse D. Kirkpatrick, Susan Su, Jaideep S. Dudani, Qian Zhong, Ahmet Bekdemir, Sangeeta N. Bhatia

**Author notes:** These authors contributed equally to this work (Ava P. Soleimany, Jesse D. Kirkpatrick).

## Abstract

Recent years have seen the emergence of conditionally activated diagnostics and therapeutics that leverage protease-cleavable peptide linkers to enhance their specificity for cancer. However, due to a lack of methods to measure and localize protease activity directly within the tissue microenvironment, the design of protease-activated agents has been necessarily empirical, yielding suboptimal results when translated to patients. To address the need for spatially resolved protease activity profiling in cancer, we developed a new class of *in situ* probes that can be applied to fresh-frozen tissue sections in a manner analogous to immunofluorescence staining. These activatable zymography probes (AZPs) detected dysregulated protease activity in human prostate cancer biopsy samples, enabling disease classification. We then leveraged AZPs within a generalizable framework to design conditional cancer diagnostics and therapeutics, and demonstrated the power of this approach in the Hi-Myc mouse model of prostate cancer, which models features of early pathogenesis. Multiplexed screening against barcoded substrates yielded a peptide, S16, that was robustly and specifically cleaved by tumor-associated metalloproteinases in the Hi-Myc model. *In situ* labeling with an AZP incorporating S16 revealed a potential role of metalloproteinase dysregulation in proliferative, pre-malignant Hi-Myc prostatic glands. Last, we incorporated S16 into an *in vivo* imaging probe that, after systemic administration, perfectly classified diseased and healthy prostates, supporting the relevance of *ex vivo* activity assays to *in vivo* translation. We envision AZPs will enable new insights into the biology of protease dysregulation in cancer and accelerate the development of conditional diagnostics and therapeutics for multiple cancer types.

## INTRODUCTION

Protease activity is dysregulated in cancer and contributes to all of its hallmarks, such as aberrant growth, neoangiogenesis, and immune evasion (1). For example, matrix metalloproteinases (MMPs) are one of several classes of enzymes known to degrade the extracellular matrix, enabling tumor invasion and metastasis (2). As a result of this dysregulation, proteases have long been considered as a potential diagnostic and therapeutic target of cancer. The past decade has seen the emergence of new classes of conditional, or “activity-based”, diagnostics (3–10) and therapeutics (11–16), which are selectively activated in response to proteases dysregulated in cancer. Probodies, for example, are a novel class of conditional therapeutics consisting of a tumor-targeting antibody masked via a protease-cleavable linker (11), which improves tumor specificity and reduces off-target toxicity. Likewise, activity-based diagnostics have demonstrated promise both pre-clinically and clinically, leveraging protease dysregulation in cancer to improve imaging specificity or intraoperative evaluation of tumor margins (4–6,17–20).

To date, conditional diagnostics and therapeutics have incorporated peptide linkers that are identified via *in vitro* screens using recombinant proteases, which do not recapitulate the complex interplay of proteases and inhibitors in native tissue contexts, nor the substantial cross-cutting inherent to short peptide sequences (1,5). As a result, the clinical utility of “conditional activation” has largely been limited to applications that do not require precise specificity for cancer (e.g. intraoperative imaging probes) or that incorporate additional layers to enhance specificity and efficacy (e.g. probodies targeting tumor-associated antigens). The development and expansion of the principle of conditional activation is currently hindered by a dearth of methods to dissect protease activity in the native tissue context (1). Methods to identify peptide substrates that are maximally cleaved by tumor-associated proteases and minimally degraded in healthy tissues would provide a modular, broadly applicable means to address this engineering challenge, and could thus facilitate the application and translation of this rapidly developing field. Specifically, multiplexed methods for substrate screening in biospecimens could yield candidate substrate linkers with high tumor specificity. Furthermore, the ability to localize these cleavage events *in situ* (i.e., in a tissue section) would enable finer dissection of cancer-associated protease biology, such as cell type-specific protease dysregulation or the distribution of active proteases across the tumor invasive front, and yield new mechanistic insights into both enzyme function and cancer biology that could in turn help validate select linkers for a given application. Finally, to accelerate translation, these methods must be compatible with clinically available biospecimens, such as biofluids, tissue microarrays, or biopsy cores.

Several methods have been developed with the aim of measuring protease activity in bulk biospecimens (21–24). Recent approaches leveraging synthetic substrates have used droplet-based microfluidics (25) or highly multiplexed peptide libraries (26,27) to profile protease activity in patient-derived biospecimens. However, these assays do not provide insight on spatial localization of protease activity within the tissue. *In situ* zymography has long been the gold standard for visualizing protease activity in tissue sections (28), but this method is limited to visualizing the activity of proteases against natural cleavage sites in gelatin and therefore lacks the modularity to accommodate the chemical diversity necessary to enable the identification or validation of disease-specific peptide substrates. Recent efforts to visualize the cleavage of synthetic peptides have relied on the expression of receptors (11,29) or integrins (8) for probe binding, limiting their modularity and generalizability. Activity-based probes (ABPs) leverage reactive warheads that covalently bind to protease active sites, enabling detection of active proteases *in vivo* and *ex vivo* (5,30). However, because ABPs detect protease activity via covalent binding rather than substrate cleavage, they are less compatible with proteases of certain catalytic classes, and *in situ* profiling with ABPs has not been used to identify peptide substrates that can be directly incorporated into activatable diagnostics and therapeutics (5).

We therefore developed modular, cleavage-based activatable zymography probes (AZPs) to localize substrate-specific proteolysis in tissue sections and applied this tool to prostate cancer (PCa) models and samples. When applied to a human tumor microarray, AZPs achieved robust classification of PCa biopsy samples, supporting their applicability to human cancer. We then leveraged a clinically relevant mouse model to demonstrate the ability of AZPs to probe the biology of cancer-associated protease dysregulation and to thus inform the bottom-up design of a conditionally activatable agent. We performed a multiplexed screen and discovered a single peptide that was specifically cleaved by disease-associated metalloproteinases in a genetically engineered mouse model (GEMM) of PCa. This substrate was then translated into an AZP, which, when applied to tissue sections, revealed metalloproteinase-dependent peptide cleavage in proliferative, pre-malignant prostatic glands. Last, we translated this substrate into a protease-activated diagnostic that selectively accumulated in diseased mouse prostates *in vivo*. These results suggest that substrates validated through *ex vivo* activity profiling can be directly translated into *in vivo* conditionally activated diagnostics or therapeutics, accelerating the design, build, and test cycle. We envision that AZPs will facilitate design of protease-activatable diagnostics and therapeutics and advance understanding of protease dysregulation in human cancer.

## RESULTS

### In situ labeling of protease activity with activatable zymography probes

We first sought to develop and validate a modular, cleavage-based zymography assay to localize protease activity within its native biological context. *In situ* labeling of protease activity offers several capabilities, including the opportunity to elucidate the spatial distribution of protease activity within a tissue (e.g., at the invasive front of a tumor) and to colocalize protease activity with histological markers (Supplementary Table S1). Drawing from previous work using cell-penetrating peptides (18,19,31), we hypothesized that peptides that consist of a cationic poly-arginine (polyR) domain complexed to an anionic poly-glutamic acid (polyE) domain via a protease-cleavable linker could enable *in situ* labeling of protease activity on tissue sections (Fig. 1a). Specifically, we hypothesized that degradation of the protease-cleavable linker component of these activatable zymography probes (AZPs) would liberate the fluorophore-labeled polyR, which could then electrostatically interact with negatively charged molecules in the tissue, enabling localization by microscopy.

**Figure 1.**
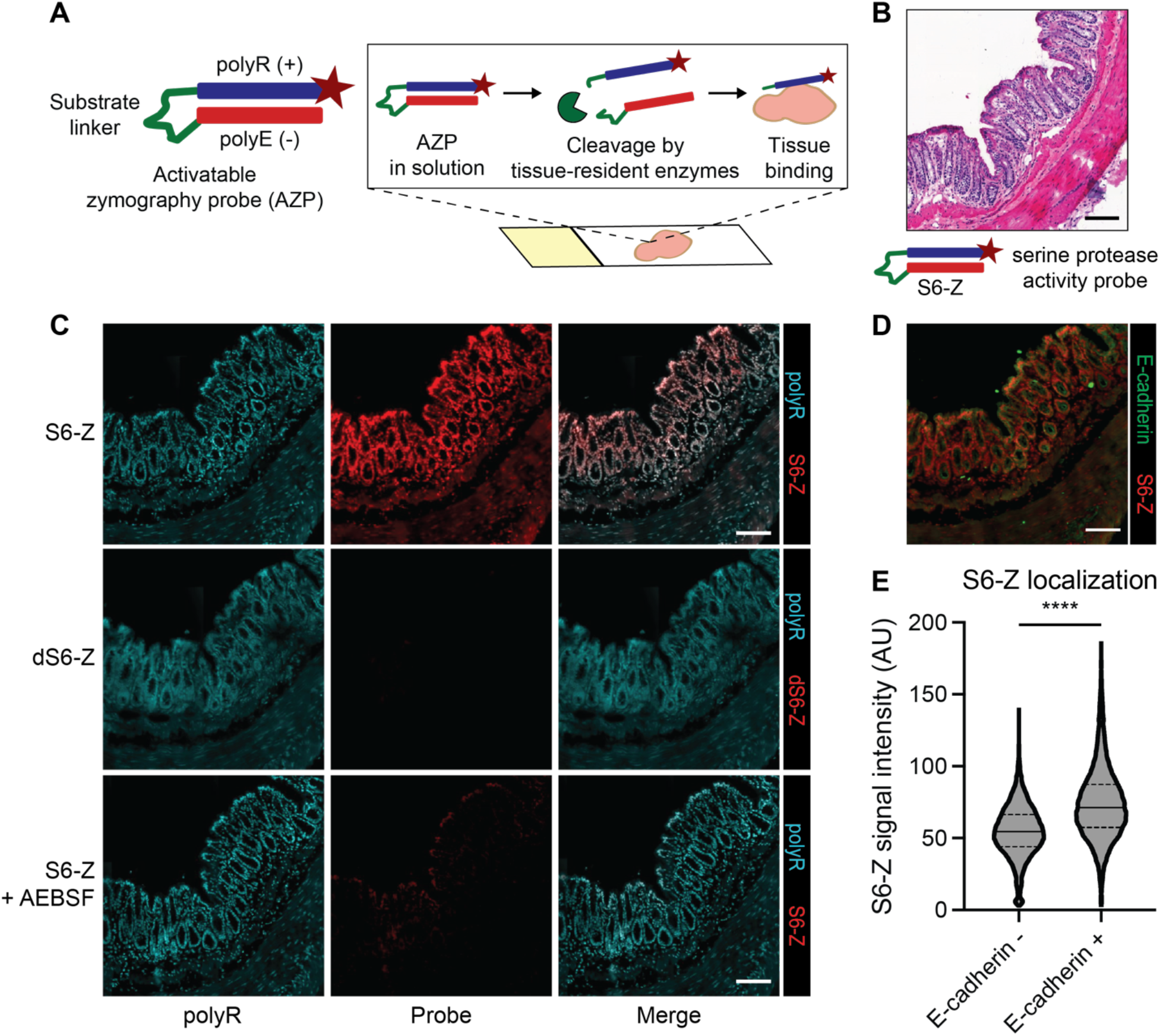
*In situ* localization of protease activity with AZPs. (a) Activatable zymography probes (AZPs). AZPs consist of a poly-arginine domain (polyR, blue) carrying a tag (e.g., fluorophore; maroon star) linked to a poly-glutamic acid domain (polyE, red) via a protease-cleavable linker (green). Upon application to frozen tissue sections, AZPs are cleaved by tissue-resident enzymes, liberating the tagged polyR which then binds the tissue. Microscopy enables *in situ* localization of substrate cleavage. (b) The serine protease-cleavable AZP, S6-Z, was applied to fresh frozen sections of healthy mouse colon. Hematoxylin and eosin (H&E) stain of a representative colon tissue region. (c) Staining of frozen colon sections with polyR (left column, teal) and either S6-Z (top, red), the uncleavable *d*-stereoisomer dS6-Z (middle, red), or S6-Z with the serine protease inhibitor AEBSF (bottom, red). (d) Region of colon tissue showing staining for S6-Z (red) and the epithelial cell marker E-cadherin (green), on a consecutive section of tissue from (c). (e) Quantification of nuclear S6-Z signal intensity from E-cadherin negative (E-cadherin-, *n* = 15864) and E-cadherin positive (E-cadherin+, *n* = 12207) cells (quantification over one representative colon section; center line represents median, dashed lines represent quartiles; two-tailed unpaired *t*-test, \*\*\*\**P* < 1 × 10^−15^). All scale bars = 100 μm.

We first tested this technique on a tissue type with a known spatial distribution of protease expression. Since colon epithelial cells secrete serine proteases (32) including urokinase plasminogen activator (uPA) (33), we synthesized a Cy5-labeled AZP, termed S6-Z, containing a substrate expected to be cleaved by serine proteases (34), and applied it to fresh-frozen sections of normal mouse colon (Fig. 1b). Fluorescence signal from S6-Z was detected in the epithelial regions of the colon, and a free polyR was used as a binding control (Fig. 1c). We synthesized a non-cleavable, *d*-stereoisomer version, termed dS6-Z, to assess whether proteolytic processing was required to activate the probe, and observed no signal following application of dS6-Z to fresh-frozen colon tissue, indicating that proteolytic cleavage was necessary for tissue binding. Furthermore, this spatially resolved S6-Z labeling was abrogated by addition of the serine protease inhibitor 4-(2-aminoethyl)benzenesulfonyl fluoride hydrochloride (AEBSF; Fig. 1c).

To characterize the extent and specificity of this activity-dependent probe localization, we applied the activatable S6-Z to colon sections and co-stained for the epithelial cell marker E-cadherin. As the primary source of serine proteases in this tissue has been shown to be colon epithelial cells (32,33), our observation that E-cadherin-positive cells exhibited increased labeling by S6-Z than E-cadherin-negative cells (*P* < 1 × 10^−15^) was consistent with our hypothesis that AZPs enable localization of cell type-specific protease activity *in situ* (Fig. 1d-e). Together, these results indicate that AZPs can directly measure peptide cleavage events *in situ* to spatially localize protease activity with low background binding.

### Discovery of human PCa-responsive protease substrates

We next set out to validate AZPs as a tool to measure enzyme activity in human tumor tissues and to thus identify peptide substrates relevant to designing conditional diagnostics and therapeutics for human cancer. To this end, we designed a library of 19 AZPs based on a panel of peptides previously found by our group to be recognized by proteases dysregulated in human PCa (34). With the exception of the protease-cleavable linker and fluorophore, these AZPs were identical in design to the serine protease-responsive S6-Z. We first sought to verify that proteolysis was required for tissue binding across the entire AZP library. We found that AZP pre-cleavage by a cognate recombinant protease with specificity for the AZP linker (MMP13 for MMP-responsive substrates; PRSS3, KLK14, or KLK2 for serine protease-responsive substrates) resulted in increased tissue binding for all 19 probes (Supplementary Fig. S1). The proteolysis-dependent tissue labeling observed across the probe library supports the modularity of the AZP platform.

Given that serine proteases are among those known to be dysregulated in human PCa(34), we selected two serine protease-responsive AZPs (S10-Z and S2-Z) to evaluate against a fresh-frozen human tissue microarray (TMA) containing biopsies from normal prostates and PCa tumors across a range of Gleason scores (Supplementary Fig. S2). Tissue binding of S10-Z (Fig. 2a, c) and S2-Z (Supplementary Fig. S3a, c) was abrogated by a broad-spectrum cocktail of protease inhibitors. We then investigated whether either of the tested AZPs preferentially labeled PCa tissue relative to normal prostate (Fig. 2b; Supplementary Fig. S3b). Both AZPs, S10-Z (*P* = 0.0063, Fig. 2d) and S2-Z (*P* = 0.0284, Supplementary Fig. S3d), exhibited significantly increased labeling of PCa relative to normal prostate tissue, with nearly all tumors demonstrating enhanced probe binding relative to controls (area under the curve (AUC) by receiver-operating characteristic; S10-Z AUC = 0.948, Fig. 2e; S2-Z AUC = 0.917, Supplementary Fig. S3e). Increased labeling of both probes was distributed across PCa tumors of all Gleason scores (Supplementary Fig. S4), suggesting that conditional diagnostics or therapeutics incorporating these linkers could be broadly applied across a heterogeneous population of patients with PCa. These findings demonstrate that serine protease activity is dysregulated in human PCa, establish the compatibility of AZPs with clinically available biospecimens such as biopsy cores, and suggest that AZPs may be used to identify peptide substrates that are preferentially cleaved by proteases in human cancer.

**Figure 2.**
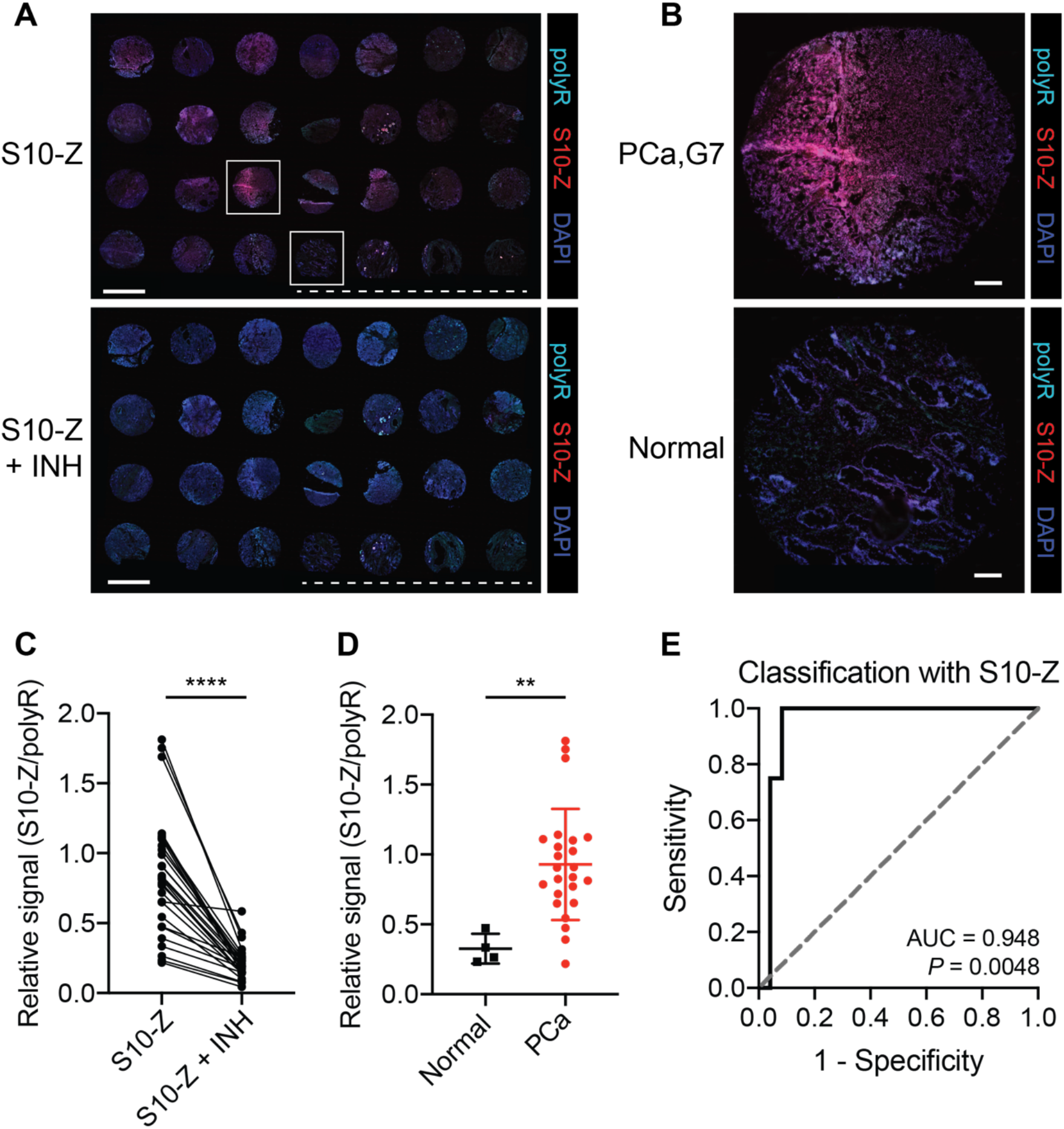
AZPs label aberrant protease activity in human PCa samples. (a) Application of S10-Z AZP (red) to a human PCa tissue microarray (TMA) consisting of 24 prostate adenocarcinoma samples and 4 normal prostate samples (S10-Z, top). A consecutive TMA was stained with S10-Z along with a cocktail of protease inhibitors (S10-Z + INH, bottom). Sections were stained with a polyR binding control (teal) and counterstained with DAPI (blue). Dotted lines are shown below normal prostate samples. Scale bars = 2 mm. (b) Higher-magnification images of boxed cores from (a) showing Gleason 7 PCa (top) and normal prostate (bottom). Scale bars = 200 μm. (c) Quantification of average S10-Z intensity relative to polyR (binding control) intensity across each TMA core (*n* = 28) for sections incubated with (S10-Z + INH) and without (S10-Z) protease inhibitors (two-tailed paired *t*-test, \*\*\*\**P* < 0.0001). (d) Quantification of relative S10-Z intensity from normal (*n* = 4) and PCa tumor (*n* = 24) cores (mean ± s.d.; two-tailed unpaired *t*-test, \*\**P* = 0.0063). (e) Receiver-operating characteristic (ROC) curve showing performance of relative AZP signal (S10-Z/polyR) in discriminating normal from PCa tumor cores (AUC = 0.948, 95% confidence interval 0.8627-1.000; *P* = 0.0048 from random classifier shown in dashed line).

### Identification of a PCa-responsive peptide substrate in the Hi-Myc model

We next aimed to incorporate our *in situ* activity probes into a generalizable pipeline to rationally design a protease-activatable diagnostic from the bottom up. We therefore turned to the Hi-Myc GEMM, wherein *c-Myc* is overexpressed in the murine prostate, resulting in invasive PCa (35). We first set out to query the landscape of proteolytic activity in Hi-Myc prostates through multiplexed substrate screening. To this end, we developed a method to measure cleavage of a multiplexed substrate panel in a single reaction volume. We selected a panel of 18 peptides previously found by our group to be recognized and cleaved by a diverse array of metallo- and serine proteases that are dysregulated in PCa (34), and appended each peptide with a unique barcode (7) consisting of the 14-mer glutamate-fibrinopeptide B (Glu-Fib) variably labeled with stable isotope-labeled amino acids. We coupled these barcoded substrates to magnetic beads such that, following incubation with a protease-containing sample and substrate cleavage, magnetic separation isolates liberated barcodes in the supernatant (Fig. 3a). Barcode concentrations are then measured by mass spectrometry, enabling quantification of substrate cleavage.

**Figure 3.**
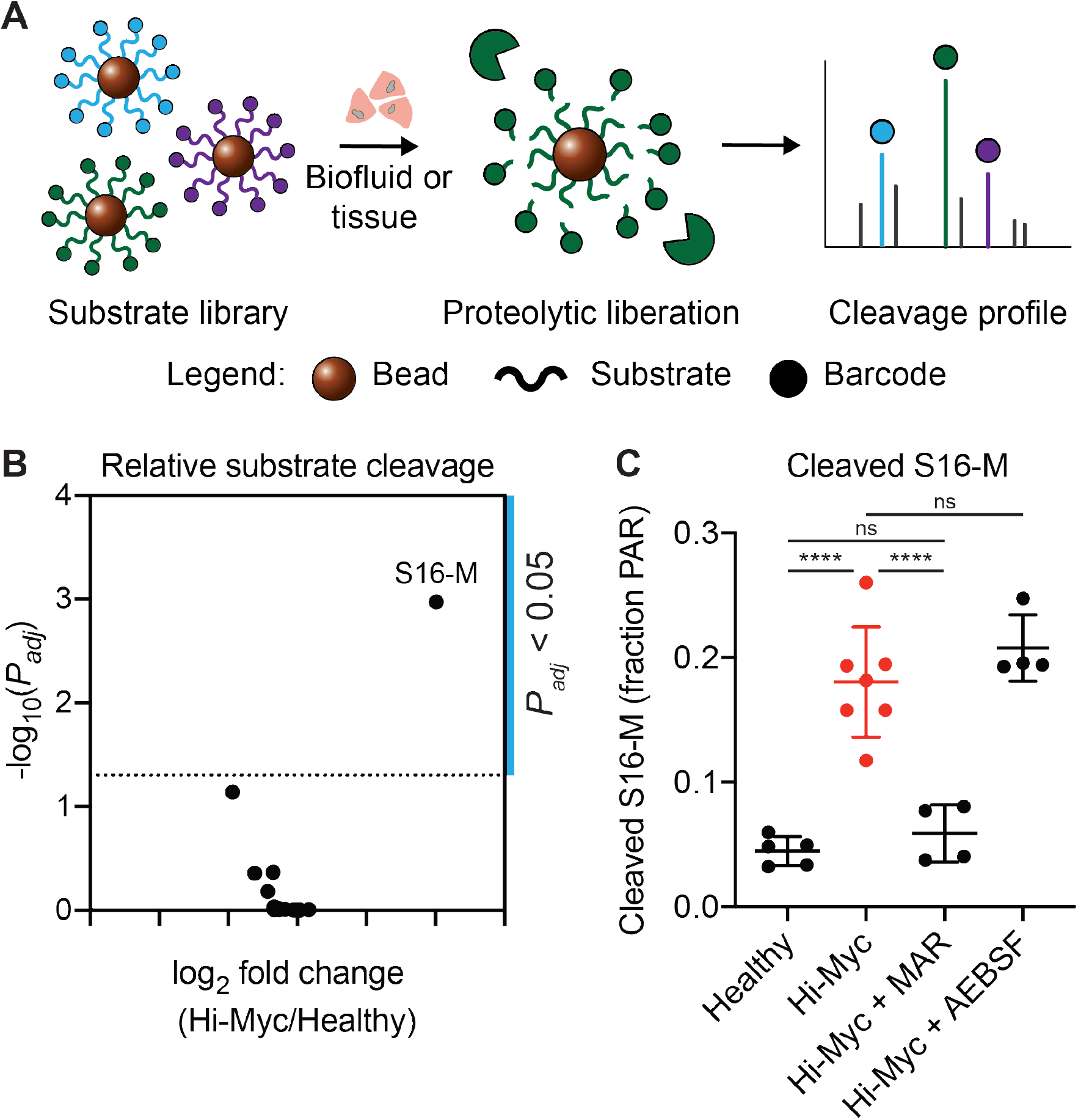
S16 is preferentially cleaved by metalloproteinases in the Hi-Myc model. (a) Multiplexed protease activity profiling with barcoded substrate libraries. Uniquely barcoded protease-cleavable peptide substrates are coupled to magnetic beads. Multiplexed bead complexes are incubated with recombinant enzymes or biospecimens. Substrate proteolysis liberates barcodes, which can be isolated via magnetic pull-down of beads and quantified to measure substrate cleavage. (b) Volcano plot showing log_2_ fold change in cleavage product concentrations between Hi-Myc (*n* = 7) and healthy (*n* = 5) prostate homogenates (x-axis) and - log_10_(*P*_*adj*_) (y-axis). Each point represents one substrate. Significance was calculated by two-tailed *t*-test followed by adjustment for multiple hypotheses using the Holm-Sidak method. Dotted line is at *P*_*adj*_ = 0.05. (c) Quantification of S16-M cleavage, measured as the fraction of total peak area ratio (PAR) signal from all screened peptides, for healthy prostate homogenates (*n* = 5) and Hi-Myc prostate homogenates, either uninhibited (Hi-Myc, *n* = 7), or in the presence of the MMP inhibitor marimastat (Hi-Myc + MAR, *n* = 4) or the serine protease inhibitor AEBSF (Hi-Myc + AEBSF, *n* = 4) (mean ± s.d.; one-way ANOVA with Tukey’s correction for multiple comparisons, \*\*\*\**P* < 0.0001, *ns* = not significant).

To validate that this assay could capture protease-substrate specificities, we incubated the peptide-functionalized bead cocktail with each of 10 recombinant proteases to assay their individual cleavage patterns and kinetics. Unsupervised hierarchical clustering of the cleavage data revealed distinct substrate specificities of the screened metallo- and serine proteases (Supplementary Fig. S5a). Furthermore, we found that the cleavage scores measured through the multiplexed bead screen generally correlated with those from a screen of quenched fluorescent versions of the same substrates incubated individually with the same selected recombinant proteases (34) (Supplementary Fig. S5b-c). Cleavage kinetics were assessed by performing magnetic separation at multiple time intervals following addition of protease and quantifying the liberated reporters by mass spectrometry (Supplementary Fig. S6). Taken together, these results indicate that the multiplexed bead assay can be used to faithfully query cleavage specificities and kinetics of target proteases against a barcoded library of peptide substrates.

We then assessed whether the multiplexed bead assay could be used to identify tumor-specific peptide substrates and patterns of protease activity dysregulation in the Hi-Myc GEMM. We screened homogenates of prostates from Hi-Myc mice and age-matched healthy controls against our multiplexed, peptide-functionalized bead library and found distinct cleavage patterns between diseased and healthy prostates (Supplementary Fig. S7a). Intriguingly, this assay revealed that a single peptide sequence (“S16”, or “S16-M” when coupled to mass barcode) was differentially cleaved by Hi-Myc prostate homogenates relative to samples from healthy control mice (Fig. 3b). Based on the results of the bead screen of peptide substrates against recombinant proteases (Supplementary Fig. S5), we predicted that the observed cleavage of the S16-M probe was likely mediated by MMP activity. Consistent with this hypothesis, we found that pre-treatment of the prostate homogenates with the MMP inhibitor marimastat (MAR) inhibited cleavage of S16-M by the Hi-Myc mouse-derived samples (*P* < 0.0001, Fig. 3c), whereas pre-treatment with the serine protease inhibitor AEBSF had no effect. Accordingly, unsupervised dimensionality reduction by principal component analysis (PCA) succeeded in separating the uninhibited homogenates (Supplementary Fig. S7b) and homogenates treated with AEBSF (Supplementary Fig. S7c), but this separation was abrogated by addition of marimastat (Supplementary Fig. S7d).

### In situ localization of protease activity in Hi-Myc tissue

The bulk, multiplexed cleavage assay identified a metalloproteinase substrate, S16, that was preferentially cleaved in prostates from Hi-Myc mice relative to healthy controls, and did not require any *a priori* knowledge of candidate protease targets. However, bulk assays inherently preclude spatial localization of the target (i.e. protease activity) within the tumor, a key factor in the design of diagnostic or therapeutic agents for cancer, such as monoclonal antibodies (36). In order to determine where S16 was cleaved within diseased prostates, we incorporated this peptide sequence into an AZP (S16-Z), verified that probe binding was protease-dependent (Supplementary Fig. S8), and applied this probe to fresh-frozen sections of prostates from Hi-Myc mice (Fig. 4a). Fluorescence signal from S16-Z was found to be broadly distributed throughout Hi-Myc tumors and was abrogated by the MMP inhibitor marimastat (Fig. 4b). In addition, we found that S16-Z labeled histologically normal glands of the Hi-Myc ventral lobe (Fig. 4a), and that this labeling was abrogated by marimastat (Supplementary Fig. S9). Furthermore, no tissue labeling was observed in sections treated with the non-cleavable probe dS16-Z, further validating that S16-Z tissue binding was proteolytically driven (Supplementary Fig. S9). Finally, a co-incubated positive control peptide (free polyR) exhibited consistent binding under all three (S16-Z, S16-Z + MAR, dS16-Z) conditions (Supplementary Fig. S9).

**Figure 4.**
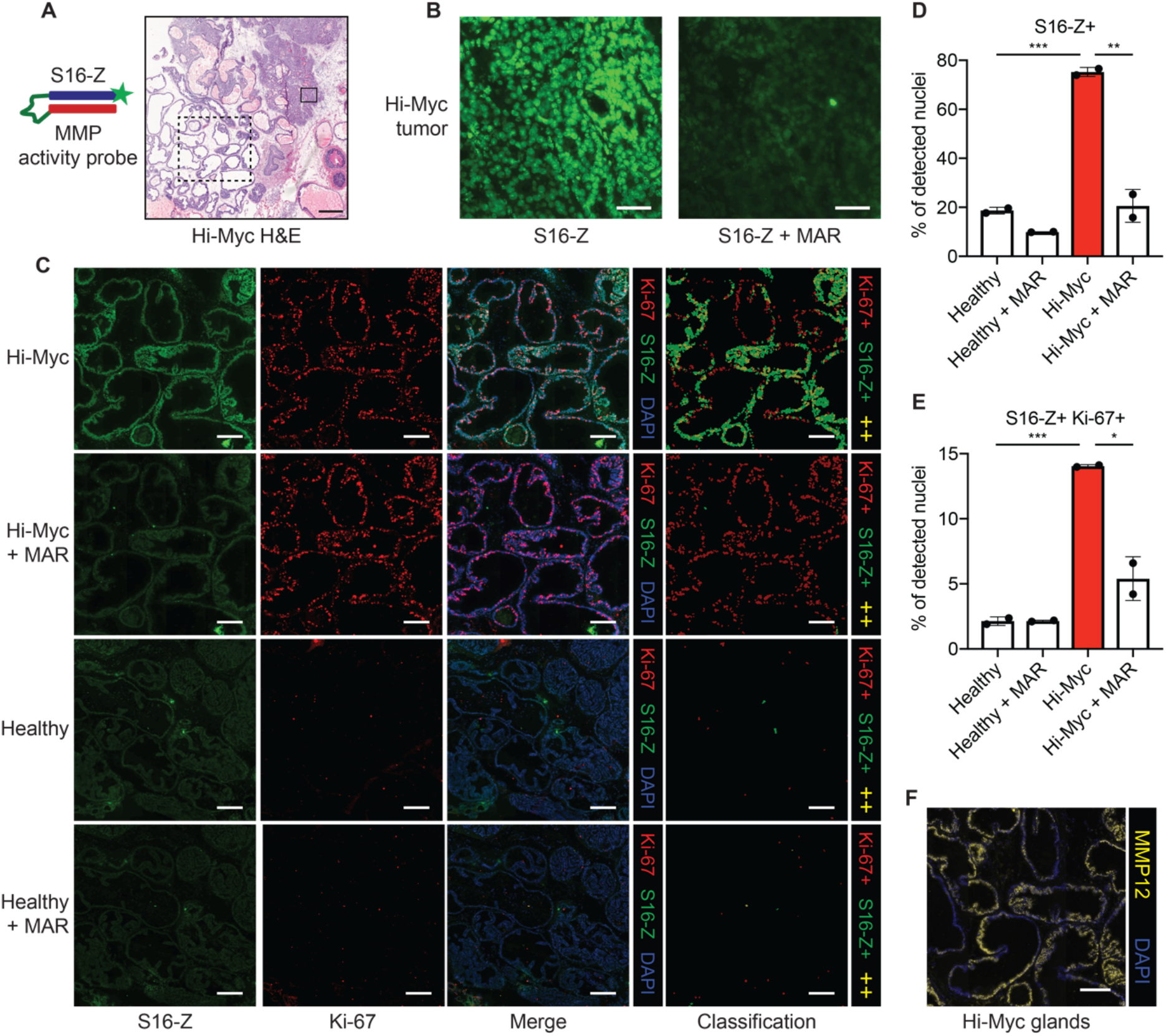
S16 cleavage localizes to established tumors and proliferative, MMP12-positive glands in the Hi-Myc model. (a) The MMP-responsive AZP S16-Z was applied to Hi-Myc prostate tissue sections. H&E staining of Hi-Myc prostate tissue, representative example shown. Solid and dashed boxes indicate tissue regions in (b) and (c), respectively. Scale bar = 500 μm. (b) Staining of boxed Hi-Myc tumor region from (a) with S16-Z (S16-Z, left). A consecutive section was stained with S16-Z in the presence of the MMP inhibitor marimastat (S16-Z + MAR, right). Scale bars = 50 μm. (c) Application of S16-Z (green), with or without MAR, to prostate tissues from Hi-Myc and healthy mice with co-staining for the proliferation marker Ki-67 (red). Sections were counterstained with DAPI (blue). Detected cells were classified on the basis of S16-Z and Ki-67 staining intensities to produce cell classification maps (red: Ki-67+; green: S16-Z+; yellow: S16-Z+ and Ki-67+, ++). Images of Hi-Myc tissues show the glandular region boxed in (a). Scale bars = 200 μm. (d) Quantification of fraction of detected nuclei positive for S16-Z in healthy, MMP-inhibited healthy (Healthy + MAR), Hi-Myc, and MMP-inhibited Hi-Myc (Hi-Myc + MAR) prostate tissue sections (*n* = 2 consecutive sections per group, >100,000 cells per section; mean ± s.d.; two-tailed unpaired *t*-test, \*\**P* = 0.00794, \*\*\**P* = 0.000803). (e) Quantification of fraction of detected nuclei positive for both S16-Z and Ki-67 (*n* = 2 consecutive sections per group, >100,000 cells per section; mean ± s.d.; two-tailed unpaired *t*-test, \**P* = 0.0185, \*\*\**P* = 0.000429). (f) Immunofluorescence staining for MMP12 (yellow) in glandular region (a) of Hi-Myc prostate tissue. Sections were counterstained with DAPI (blue). Scale bar = 200 μm.

Though these glands appeared histologically normal, their observed metalloproteinase activity, which is associated with several cancer hallmarks (1), raised the possibility of early-stage malignant transformation in these regions. In an effort to undertake further biological characterization, we applied S16-Z to prostate tissue sections from Hi-Myc and age-matched healthy mice and co-stained for the proliferation marker Ki-67 (37). Strikingly, we observed robust Ki-67 staining in S16-Z-positive, but otherwise histologically normal, glands of the Hi-Myc prostate, but minimal Ki-67 staining and reduced S16-Z labeling in a histologically similar region of ventral prostate from healthy mice (Fig. 4c). Segmentation-based, cell-level classification indicated the presence of cells that stained positively for both S16-Z and Ki-67 within this glandular region of Hi-Myc tissue, but an absence of such double-positive cells in healthy prostatic glands (Fig. 4c). Similar analyses of a Hi-Myc tumor showed the presence of S16-Z-positive tumor cells that also stained positively for Ki-67 (Supplementary Fig. S10). Furthermore, S16-Z labeling across both proliferative glands and established tumors in Hi-Myc prostates was abrogated by marimastat, while MMP inhibition appeared to have a minimal effect on S16-Z staining in healthy tissues (Fig. 4c; Supplementary Fig. S10).

To quantify these findings, we performed cell-by-cell quantification of S16-Z and Ki-67 staining intensities across entire Hi-Myc and healthy prostate tissue sections (Supplementary Fig. S11; Supplementary Fig. S12). This analysis indicated that Hi-Myc tissue exhibited a significantly increased proportion of S16-Z-positive cells relative to healthy prostates (*P* = 0.000803, Fig. 4d). Furthermore, incubation with marimastat significantly reduced the number of S16-Z-positive cells in Hi-Myc prostates (*P* = 0.00794, 72.7% decrease; Fig. 4d). Finally, as suggested by cell-based segmentation analysis, we observed a significantly greater fraction of S16-Z, Ki-67 double-positive cells in Hi-Myc prostate sections relative to healthy tissues (*P* = 0.000429, Fig. 4e). Taken together, these *in situ* measurements and quantitative analyses revealed significant dysregulation of MMP activity and cell proliferation in both established tumors and in histologically normal glands of Hi-Myc tissue that was absent in healthy prostates.

Motivated by these results, we next sought to nominate candidate metalloproteinases that could be responsible for S16-M and S16-Z cleavage in the Hi-Myc model. We first screened a quenched fluorescent version of this substrate, S16-Q, against a panel of recombinant MMPs and found that this probe was broadly MMP-cleavable (Supplementary Fig. S13a-b). Next, we queried previously reported gene expression data from the Hi-Myc model (35) and found that MMP12, which efficiently cleaved S16-Q *in vitro*, was significantly upregulated at the gene expression level in Hi-Myc PCa relative to healthy prostate tissue (Supplementary Fig. S13c). We then validated this transcript at the protein level and found a significant increase in MMP12 in Hi-Myc prostates relative to age-matched healthy controls (*P* = 0.0034, Supplementary Fig. S13d). Additionally, Hi-Myc prostates stained positively for MMP12 while healthy prostates did not (Supplementary Fig. S13e). MMP12 was also found to localize broadly throughout the histologically normal, S16-Z, Ki-67-positive glands of Hi-Myc tissue (Fig. 4f). Overall, these results suggest that MMP12 may be responsible for S16 cleavage in Hi-Myc prostates, highlighting the power of AZPs to enable *in situ* localization and biological characterization of protease dysregulation in cancer.

### In vivo validation of a protease-activated probe in the Hi-Myc model

Having leveraged a multiplexed screen and AZP profiling to identify and localize S16 cleavage in Hi-Myc prostates, we sought to rationally design an *in vivo* probe that could be specifically activated within diseased prostates. To this end, we synthesized a quenched imaging probe, S16-QZ, that incorporated S16 as a protease-cleavable linker. In this design, degradation of the protease-cleavable linker liberates the fluorophore-tagged polyR, resulting in a positively charged, tissue-binding peptide with Cy5 fluorescence. Accordingly, *in vitro* incubation of S16-QZ with MMP12 resulted in fluorescence activation, whereas the uncleavable dS16-QZ remained quenched even after incubation with the protease (Fig. 5a). To verify that tissue binding was protease-dependent, we pre-cleaved S16-QZ with recombinant MMP12, incubated the probe with tissue sections at 4° C to allow tissue binding in the absence of protease activity, and observed significantly enhanced tissue fluorescence relative to intact S16-QZ (Fig. 5b, c).

**Figure 5.**
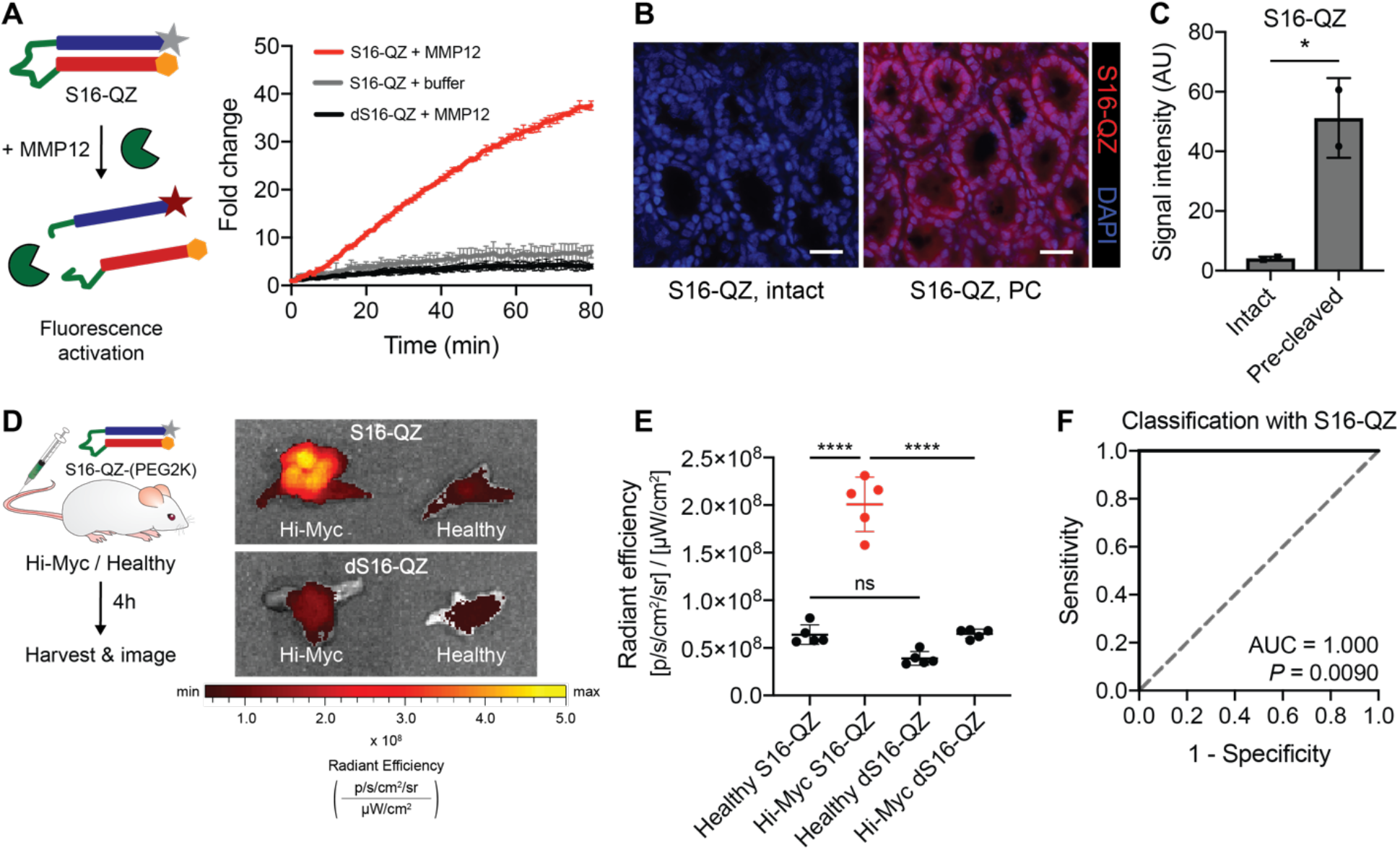
An activatable imaging probe incorporating S16 enables classification of prostate cancer *in vivo*. (a) The quenched probe S16-QZ, consisting of a fluorophore-tagged polyR (grey star + blue rectangle) and quencher-tagged polyE (orange hexagon + red rectangle) linked via an MMP-cleavable linker (green), was incubated with recombinant MMP12, and fluorescence activation was monitored over time (left). Dequenching measurements for S16-QZ and *d*-stereoisomer dS16-QZ following incubation with recombinant MMP12 (right; *n* = 3 replicates per condition; mean ± s.d.). (b) Binding of intact or MMP12 pre-cleaved (PC) S16-QZ to fresh frozen sections of healthy mouse colon tissue following incubation at 4° C. Scale bars = 25 μm. (c) Quantification of S16-QZ binding for intact probe or probe pre-cleaved by MMP12 (*n* = 2 replicates; mean ± s.d.; two-tailed unpaired *t*-test, *\*P* = 0.0382). (d) S16-QZ-(PEG2K) or uncleavable dS16-QZ-(PEG2K) was administered intravenously into age-matched Hi-Myc and healthy mice. 4 hours after injection, prostates were excised and imaged by IVIS (*n* = 5 per group). Representative prostates shown at right. (e) Quantification of epifluorescence radiant efficiency from experiment in (d) via IVIS imaging (*n* = 5 per group; mean ± s.d.; one-way ANOVA with Tukey’s correction for multiple comparisons, \*\*\*\**P* < 0.0001, ^ns^*P* = 0.0990). (f) Receiver-operating characteristic (ROC) curve showing performance of S16-QZ-(PEG2K) signal in discriminating healthy from Hi-Myc prostate explants (AUC = 1.000, 95% confidence interval 1.000-1.000; *P* = 0.0090 from random classifier shown in dashed line).

Having verified that proteolytic processing was required for both fluorescence activation and tissue binding of this PCa-targeted probe, we next sought to assess whether this probe could enable detection of disease after systemic *in vivo* administration. S16-QZ and the uncleavable control peptide, dS16-QZ, were PEGylated to improve solubility, increase circulation time, and drive tissue accumulation (31), and were subsequently administered intravenously into Hi-Myc mice and age-matched healthy controls (Fig. 5d). Mice were assessed at 32-36 weeks of age, at which point adenocarcinoma is typically present in Hi-Myc prostates (35). Significantly increased Cy5 fluorescence was observed in explanted prostates of Hi-Myc mice injected with S16-QZ relative to the uncleavable dS16-QZ (*P* < 0.0001, Fig. 5e), whereas no such difference between the probes was observed in healthy prostates (*P* = 0.0990, Fig. 5e). Furthermore, Cy5 fluorescence after S16-QZ injection was found to be significantly increased in Hi-Myc prostates relative to healthy prostates (*P* < 0.0001, Fig. 5e), enabling perfect discrimination of Hi-Myc and healthy mice (AUC = 1.000, Fig. 5f). These results demonstrate that peptide substrates nominated via *ex vivo* activity profiling in tumor specimens can be directly incorporated into conditionally activated agents for *in vivo* tumor targeting and detection.

## DISCUSSION

In this work, we developed a modular assay to localize protease activity *in situ* and incorporated this method into a generalizable pipeline to rationally design protease-activated diagnostics and therapeutics. By applying our localization probes, AZPs, to a human PCa tumor microarray, we identified two peptides that are preferentially cleaved by human PCa-associated proteases, thereby supporting the ability of AZPs to directly validate candidate cleavable linkers in patient-derived tissue samples, prior to *in vivo* administration. Next, we demonstrated a bottom-up approach to the development of activatable diagnostics in the Hi-Myc PCa GEMM. We identified a peptide, S16, cleaved specifically by PCa-associated metalloproteinases, and found, using AZPs, that its cleavage localized broadly throughout tumors and proliferative, MMP12-positive glands in diseased prostates. Finally, we demonstrated that this disease-responsive substrate characterized via *ex vivo* activity profiling could be integrated into a protease-activated diagnostic for *in vivo* use.

Integrating multiplexed screens that nominate lead substrates with cleavage-based zymography assays that validate candidates *in situ* presents a new framework relevant to both discovery and engineering efforts that seek to understand and leverage protease dysregulation in cancer (Fig. 6). Each of these methods provides key advantages—multiplexed screening enables nomination of candidate substrates without any prior knowledge of enzyme targets, but *in situ* labeling is necessary to explore underlying biology and to validate the relevance of these substrates for specific applications. For example, an activatable therapeutic like a probody would ideally incorporate a peptide substrate that is cleaved broadly throughout the tumor, whereas an activatable imaging probe designed to monitor responses to immunotherapy might leverage a peptide that is strictly cleaved by proteases produced by tumor-infiltrating lymphocytes.

**Figure 6.**
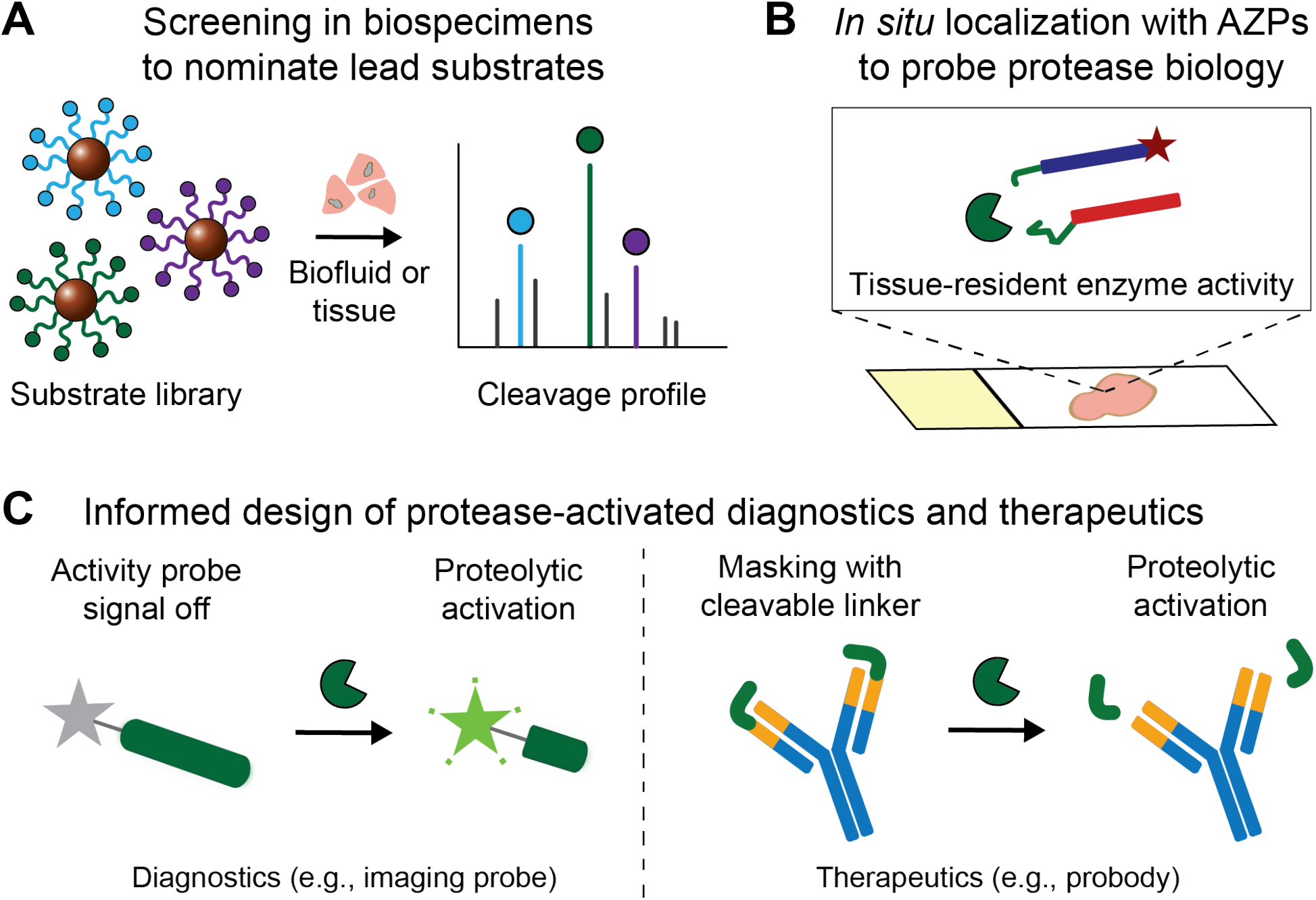
Pipeline to design conditional cancer diagnostics and therapeutics. (a) Multiplexed protease activity profiling with barcoded substrates can be used to nominate lead peptides preferentially cleaved in diseased tissues. (b) Activatable zymography probes (AZPs) localize protease activity in fresh-frozen tissue sections, validating the relevance of substrate candidates for a given application and further dissecting the biology of protease dysregulation in cancer. (c) Disease-responsive peptide substrates nominated from *ex vivo* activity profiling can be directly incorporated into protease-activated diagnostics, e.g., *in vivo* imaging probes, or therapeutics, e.g., probodies, proteolytically activated antibodies that require linker cleavage for target binding, for *in vivo* evaluation.

Here, we demonstrate the power of this approach in the Hi-Myc model of PCa, identifying the disease-responsive peptide substrate S16 from a multiplexed screen and using *in situ* activity profiling with AZPs, coupled with immunostaining and quantitative analysis, to find that S16 cleavage localized to both tumor cells and Ki-67-positive glands that had not yet progressed to adenocarcinoma. This ability to quantitatively co-localize protease activity signals with other markers may enable concurrent identification of both disease-specific peptide linkers and biological targets for rational design of protease-activated diagnostics and therapeutics. Last, we show that an *in vivo* imaging probe incorporating S16 exhibited enhanced signal in Hi-Myc prostate explants, establishing the relevance of *ex vivo* activity profiling for *in vivo* translation. Together with this *in vivo* validation, the observation that AZPs localized S16 cleavage in pre-malignant prostatic glands suggests that, if incorporated into a diagnostic or therapeutic, this substrate could enable detection or interception of early-stage disease in the Hi-Myc model.

AZPs offer several advantages over existing methods of measuring and localizing protease activity *in situ*. While *in situ* activity profiling with fluorescently tagged probodies has recently been shown to correlate with *in vivo* therapeutic efficacy, this approach requires target antigen expression for probe binding and is therefore limited in its generalizability (29). AZPs address this limitation in that they provide a modular, antigen-independent means to directly measure substrate-specific protease activity. Activity-based probes, which covalently bind to protease active sites, consist of two components (a recognition sequence and a reactive functional group) that must be redesigned for each new protease target (5). AZPs, in contrast, consist of two constant regions (polyR and polyE, Fig. 1b) and a single protease-cleavable linker that can be replaced with any sequence of interest. This modularity enabled us to synthesize a library of 19 AZPs, all of which were dependent on proteolysis for tissue labeling (Supplementary Fig. S1). Therefore, we expect that AZPs could readily be designed to incorporate any candidate peptide substrate, a degree of modularity unmatched by existing *in situ* probes.

This work also establishes AZPs as a molecular tool to probe protease-mediated biological processes *in situ* and to thereby functionally dissect the crucial roles of proteases in cancer development and progression. Many proteases, including MMPs, are synthesized as zymogens and require proteolytic processing for activation (2). Furthermore, protease activity is regulated by endogenous protease inhibitors, including members of the tissue inhibitor of metalloproteinases (TIMP) and serpin families (5). Because proteases exert their function through their activity, direct measurements of protease activity rather than abundance may enable new insights into the roles of enzymatic biology in cancer. In this work, we uncovered an increase in metalloproteinase activity in established prostate tumors and, unexpectedly, in histologically normal prostatic glands in the Hi-Myc model. This finding was powered by the identification of S16, a metalloproteinase-sensitive peptide cleaved by proteases in the Hi-Myc model, followed by *in situ* labeling with S16-Z. Additional investigation revealed that S16-Z-positive, histologically normal glands, in addition to established tumors, were also positive for the proliferation marker Ki-67 and overexpressed MMP12. MMP12 has previously been shown to inhibit tumor growth in mice (38), an effect that appears to derive from its ability to cleave plasminogen and generate angiostatin, a potent angiogenesis antagonist (39). Though mechanistic experiments will be necessary to fully elucidate the role of MMP12 in the pathogenesis of PCa, the intensity of S16-Z staining of Ki-67-positive glands in the Hi-Myc prostate raises the prospect of a robust, MMP12-mediated anti-angiogenic response that parallels and counterbalances dysregulated cell division in this disease model. As demonstrated by these data, integrating *in situ* protease activity profiling, immunostaining, and quantitative image analysis could provide a powerful paradigm to dissect enzymatic biology in cancer initiation, progression, and spread.

Though the cleavage-based AZPs presented here offer several advantages over existing probes, their primary limitation is specificity, as proteases of the same class exhibit substantial cross-cutting against short peptides. Multiplexed screens and *in situ* labeling with AZPs can be readily deployed as tools to nominate and validate peptide linkers for conditionally activated diagnostics or therapeutics, which must be selective for cancer relative to healthy tissue but need not be specific to any one protease (11). In contrast, further optimization is needed before such cleavage-based methods can be reliably leveraged as a tool to dissect protease biology with single enzyme specificity. High-throughput screens of phage-displayed substrate libraries (40) or peptides enriched with unnatural amino acids (41,42) may yield protease substrates with dramatically improved specificities.

In summary, we have engineered a new, highly modular class of *in situ* activity probes, AZPs, to identify tumor-specific peptide substrates and to localize their cleavage in tissue sections. We integrated multiplexed substrate screening with AZP profiling to achieve the bottom-up design of an activatable imaging probe in a PCa GEMM and also showed that AZPs can be used to directly probe protease activity in human biopsy specimens. We expect that AZPs, which can be readily adapted to any tumor type or set of peptide substrates, will help elucidate the multifaceted roles of proteases in tumor biology and accelerate the development of next-generation agents that leverage protease activity to detect and target cancer.

## MATERIALS AND METHODS

### Peptide synthesis

All peptides were synthesized by CPC Scientific (Sunnyvale, CA) and reconstituted in dimethylformamide (DMF) unless otherwise specified.

### Hi-Myc PCa model

All animal studies were approved by the Massachusetts Institute of Technology (MIT) committee on animal care (protocol 0420-023-23). Reporting complied with Animal Research: Reporting In Vivo Experiments (ARRIVE) guidelines. Hi-Myc mice (FVB-Tg(ARR2/Pbsn-MYC)7Key/Nci) were obtained from Jackson Labs. The Hi-Myc model was generated as previously described (35). Age-matched healthy male FVB/NJ mice (Jackson Labs) were used as controls in all experiments.

### Mouse dissections and tissue isolation

For prostate tissue, urogenital organs from Hi-Myc mice and age-matched healthy male FVB/NJ mice (Jackson Labs) were isolated, and prostates were microdissected. Adipose tissue surrounding the prostates was removed using forceps, and the entirety of the mouse prostate (ventral, dorsal, lateral, and anterior lobes) was separated from the urethra. For preparation for tissue homogenization, prostates were flash frozen in liquid nitrogen and stored at −80° C until use. For preparation for cryosectioning, prostates were immediately embedded in optimal-cutting-temperature (OCT) compound (Sakura), and frozen using isopentane chilled with dry ice. Colons were harvested from healthy C57BL/6J mice (Jackson Labs), rinsed with PBS to remove feces, filled with undiluted OCT using a blunt 20-gauge needle, embedded in OCT, and frozen using isopentane chilled with dry ice.

### Synthesis of peptide-functionalized magnetic beads

For synthesis of peptide-functionalized magnetic beads, amine-functionalized magnetic beads (Dynabeads™ M-270 Amine, Invitrogen) were washed and resuspended in PBS prior to coupling, per manufacturer’s instructions. For conjugation of peptides to magnetic beads, the beads were first reacted with MAL-PEG(5k)-SVA (Laysan Bio) in PBS at room temperature for 30 minutes with gentle rotation to introduce sulfhydryl-reactive maleimide handles. Cysteine-terminated peptides (CPC Scientific) with mass-encoded barcodes were then reacted in PBS with up to 30% DMF to the PEG-coated magnetic beads overnight at room temperature with gentle rotation. Following overnight incubation, any remaining sulfhydryl-reactive handles were quenched with L-cysteine (Sigma Aldrich) by reacting for at least 15 minutes at room temperature. Peptide-functionalized magnetic beads were then pooled, washed in bead wash buffer (0.5% (w/v) bovine serum albumin (BSA), 0.01% (v/v) Tween-20 in PBS) according to manufacturer’s instructions, buffer exchanged into PBS, and then resuspended into PBS at the original bead concentration provided by the manufacturer. Peptide-functionalized magnetics beads were stored at 4° C until use. Relative substrate concentrations were quantified via LC-MS/MS analysis of stock bead cocktail solution.

### LC-MS/MS reporter quantification

LC-MS/MS was performed by Syneos Health (Princeton, NJ) using a Sciex 6500 triple quadrupole mass spectrometer. Samples were treated with ultraviolet irradiation to photocleave the 3-amino-3-(2-nitro-phenyl)propionic acid (ANP) and liberate the barcoded reporter from residual peptide fragments. Samples were extracted by solid-phase extraction and analyzed by multiple reaction monitoring by LC-MS/MS to quantify concentrations of each barcoded mass variant. Analyte quantities were normalized to a spiked-in internal standard, and concentrations were calculated from a standard curve using peak area ratio (PAR) to the internal standard. Initial peptide concentrations were quantified via LC-MS/MS analysis of stock bead solution used in the cleavage reaction.

### In vitro recombinant protease cleavage assays

For bead cleavage assays, magnetic beads functionalized with mass-barcoded peptide substrates were synthesized and prepared as described above. Recombinant proteases were purchased from Enzo Life Sciences, R&D Systems, and Haematologic Technologies. Assays for each recombinant protease were conducted in appropriate enzyme buffer according to manufacturer specifications, with 12.5 nM recombinant enzyme. For each cleavage reaction, 1.2 × 10^8^ beads, consisting of an isokinetic mixture of beads carrying unique substrate-reporter pairs and resuspended to a volume of 60 μL in appropriate enzyme buffer, were added to a PCR reaction tube, and 20 μL of recombinant protease was added and mixed with the bead solution to achieve a final enzyme concentration of 12.5 nM in 80 μL reaction volume. Reactions were incubated at 37° C with gentle rotation (to prevent bead settling) for 1 hour. Following incubation and proteolysis, two rounds of magnetic bead pull-down were used to isolate liberated reporters present in the supernatant. Samples were frozen at −80° C until analysis by LC-MS/MS as described above. Initial peptide concentrations were quantified via LC-MS/MS analysis of stock bead solution used in the cleavage reaction. To determine the relative concentrations of cleavage-liberated reporters, liberated reporter PAR values were normalized to stock reporter PAR values.

For fluorescence cleavage assays, intramolecularly quenched peptide substrates (S16-Q, S16-QZ, dS16-QZ) were reacted with recombinant proteases to characterize cleavage kinetics. Substrates were incubated with recombinant MMP12 (Enzo Life Sciences) at a final peptide concentration of 1 μM and a final enzyme concentration of 12.5 nM in a reaction volume of 30 μL in MMP-specific buffer (50 mM Tris, 300 mM NaCl, 10 mM CaCl_2_, 2 μM ZnCl_2_, 0.02% (v/v) Brij-35, 1% (w/v) BSA, pH 7.5) at 37° C. Proteolytic cleavage of substrates was quantified by increases in fluorescence over time as measured by fluorimeter (Tecan Infinite M200 Pro).

### Bead cleavage assays with tissue homogenates

Prostates from Hi-Myc and age-matched healthy FVB/NJ male mice were harvested and fresh frozen as described above. Prior to fluorogenic cleavage assay, prostates were placed in tubes with high-impact zirconium beads (Benchmark Scientific), and PBS was added to achieve a concentration of 100 to 300 mg tissue per mL of PBS. Homogenization was performed using a BeadBeater homogenizer (BioSpec). Bradford assay (Bio-Rad) was performed, and all samples were prepared to twice (~2 mg/mL) the desired final assay protein concentration (~1 mg/mL). For inhibited conditions, 1 mM AEBSF (Sigma Aldrich) or 2 mM marimastat (Sigma Aldrich) were pre-incubated with tissue homogenates for at least 30 minutes prior to cleavage assay.

Magnetic beads functionalized with mass-barcoded peptide substrates were synthesized and prepared as described above. For each cleavage reaction, 1.2 × 10^8^ beads, comprised of a multiplexed cocktail of beads carrying unique substrate-reporter pairs and resuspended to a volume of 40 μL in PBS, were added to a PCR reaction tube, and 40 μL of tissue sample (2 mg/mL protein concentration) was added and mixed with the bead solution to achieve a final protein concentration of 1 mg/mL in 80 μL reaction volume. Reactions were incubated at 37° C with gentle rotation (to prevent bead settling) for 1 hour. Following incubation and proteolysis, two rounds of magnetic separation were used to isolate liberated reporters. Samples were frozen at −80° C until analysis by LC-MS/MS as described above. To determine the relative concentrations of cleavage-liberated reporters, reporter PAR values were scaled by the per-sample sum of PARs across all reporters, to account for tissue-to-tissue differences in substrate and protein concentration.

### In situ zymography with AZPs

Organs were extracted and immediately embedded and frozen in optimal-cutting-temperature (OCT) compound (Sakura). Cryosectioning was performed at the Koch Institute Histology Core. Prior to staining, slides were air dried and fixed in ice-cold acetone for 10 minutes. After hydration in PBS (3×5 minutes), tissue sections were blocked in protease assay buffer (S6-Z: 50 mM Tris, 0.01% (v/v) Tween 20, 1% (w/v) BSA, pH 7.4; S2-Z/S10-Z: 50 mM Tris, 150 mM NaCl, 10 mM CaCl_2_, 0.05% (v/v) Brij-35, 1% (w/v) BSA, pH 7.5; S16-Z: 50 mM Tris, 300 mM NaCl, 10 mM CaCl_2_, 2 μM ZnCl_2_, 0.02% (v/v) Brij-35, 1% (w/v) BSA, pH 7.5) for 30 minutes at room temperature. Blocking buffer was aspirated, and solution containing fluorescently labeled AZP (1 μM) and a free poly-arginine control (polyR, 0.1 μM) diluted in the relevant assay buffer was applied. Slides were incubated in a humidified chamber at 37° C for 4 hours. For inhibited controls, 400 μM AEBSF (Sigma Aldrich) or 1 mM marimastat (Sigma Aldrich) was added to the buffer at both the blocking and cleavage assay steps for S6-Z and S16-Z experiments, respectively. For co-staining experiments, primary antibodies (E-cadherin, AF648, R&D Systems, 20 μg/mL; Ki-67, ab15580, Abcam, 1 μg/mL) were included in the AZP solution. Following AZP incubation, slides were washed in PBS (3×5 minutes), stained with Hoechst (5 μg/mL, Invitrogen) and the appropriate secondary antibody if relevant (Invtirogen), washed in PBS (3×5 minutes), and mounted with ProLong Diamond Antifade Mountant (Invitrogen). Slides were scanned on a Pannoramic 250 Flash III whole slide scanner (3DHistech).

### AZP library characterization

AZPs (10 μM final concentration) were incubated with recombinant protease (100 nM final concentration) in 60 μL final volume of appropriate enzyme buffer at 37° C for at least 2 hours to run cleavage reactions to completion. Recombinant proteases were selected on the basis of substrate characterization and cleavage specificity (Fig. 2). S1-Z, S3-Z, S7-Z, S14-Z, S15-Z, S16-Z, and S19-Z were incubated with recombinant MMP13 (Enzo Life Sciences); S2-Z, S6-Z, S9-Z, S11-Z, S12-Z, S13-Z, S17-Z, and S20-Z AZPs were incubated with recombinant PRSS3 (R&D Systems); S5-Z, S8-Z, and S18-Z were incubated with recombinant KLK14 (R&D Systems); and S10-Z was incubated with recombinant KLK2 (R&D Systems). Following pre-cleavage with recombinant enzymes, AZP solutions were diluted to a final peptide concentration of 1 μM in appropriate enzyme buffer. Cognate in-tact AZPs (1 μM in appropriate enzyme buffer) and pre-cleaved AZPs were applied to fresh frozen sections of normal mouse colon (slide preparation previously described) and incubated at 37° C for 30 minutes. Following AZP incubation, slides were washed, stained with Hoechst, mounted, and scanned as previously described. The MMP12-responsive probes incorporating the substrate S16, S16-Z and S16-QZ, were additionally characterized using pre-cleavage by MMP12 (Enzo Life Sciences), followed by application to fresh frozen sections of normal mouse colon and incubation at 4° C for 30 minutes.

### AZP analysis of human PCa tissue microarray (TMA)

Fresh frozen human PCa TMAs (BioChain Institute, Inc.; T6235201) were air dried and fixed in ice-cold acetone for 10 minutes. After hydration in PBS (as previously described), tissue sections were blocked in serine protease assay buffer (50 mM Tris, 150 mM NaCl, 10 mM CaCl_2_, 0.05% (v/v) Brij-35, 1% (w/v) BSA, pH 7.5) for 30 minutes at room temperature. Blocking buffer was aspirated, and solution containing AZP (1 μM) and a free poly-arginine control (polyR, 0.1 μM) diluted in assay buffer was applied. Slides were incubated in a humidified chamber at 37° C for 4 hours. For inhibited controls, protease inhibitor cocktail (P8340, Sigma Aldrich) spiked with 400 μM AEBSF (Sigma Aldrich) and 1 mM marimastat (Sigma Aldrich) was added to the buffer at both the blocking and cleavage assay steps. Following AZP incubation, slides were washed, stained with Hoechst, mounted, and scanned as previously described.

### ELISA for MMP12

Tissue levels of MMP12 in Hi-Myc and healthy FVB/NJ prostate homogenates were measured by ELISA according to manufacturer’s protocol (ab246540, abcam). Prostate homogenates were prepared as previously described. Protein concentration was measured via Bradford assay (Bio-Rad) and normalized across samples.

### Immunofluorescence staining for MMP12

Slides of fresh-frozen tissue sections were prepared as described above. Slides were blocked in serum-based blocking buffer and incubated with primary antibody against MMP12 (MA5-32011, Invitrogen, 1:50, 20 μg/mL). Slides were washed in PBS (3×5 minutes), incubated with the appropriate secondary antibody (Invitrogen) and Hoechst (5 μg/mL, Invitrogen), and washed in PBS (3×5 minutes). Slides were mounted and scanned as previously described.

### Quantification of AZP and immunofluorescence staining

AZP and immunofluorescence staining was quantified in QuPath 0.2.0-m9 (43). To perform cell-by-cell analysis, segmentation was performed via watershed cell detection on the DAPI (nuclear) channel. Binary classification was performed to identify cells that were positive or negative for E-cadherin, S16-Z, and Ki-67. E-cadherin positive cells had maximum cytoplasmic E-cadherin intensity of greater than or equal to 65 AU. S16-Z positive cells had mean nuclear S16-Z intensity of greater than or equal to 20 AU. Ki-67 positive cells had mean nuclear Ki-67 intensity of greater than or equal to 10 AU. All cells that were not positive for a particular marker were considered negative for that marker. Classification maps were generated using the object classification function in QuPath. Density plots were generated using the dscatter function in MATLAB R2017b. Additional statistical analyses were conducted in GraphPad 7.0 (Prism).

To quantify relative binding of pre-cleaved and intact AZPs, mean nuclear AZP staining was averaged across all cells in each section. For TMA analysis, AZP staining was calculated as a fold change of the mean nuclear AZP signal over the mean nuclear polyR signal. All nuclei were averaged across the entire TMA core. Nuclei with a polyR intensity of less than 3 were excluded from analysis.

### Synthesis of activatable probes for in vivo administration

The peptides S16-QZ and dS16-QZ were reconstituted to 1 mg/mL in water. Peptides were then reacted with mPEG-Maleimide, MW 2000 g/mol (Laysan Bio), for PEG coupling via maleimide-thiol chemistry. After completion of the reaction, the final compounds were purified using HPLC. All reactions were monitored using HPLC connected with mass spectrometry (LC/MS). Characterization of final compounds, S16-QZ-(PEG2K) and dS16-QZ-(PEG2K), using HPLC and MALDI-MS indicated that products were obtained with more than 90% purity and at the expected molecular weight.

### In vivo administration of activatable probes and imaging of explanted prostate tissues

Male Hi-Myc mice (FVB-Tg(ARR2/Pbsn-MYC)7Key/Nci; 32-36 wks) and age-matched FVB/NJ healthy controls (Jackson Labs; 28-36 wks) were anesthetized using isoflurane inhalation (Zoetis). S16-QZ-(PEG2K) or dS16-QZ-(PEG2K) (4.5 nmoles in 0.9% NaCl) was administered intravenously via tail-vein injection. Four hours after probe injection, mice were euthanized by isoflurane overdose followed by cervical dislocation, and prostates were dissected as previously described. Explanted prostates were imaged on an *in vivo* imaging system (IVIS, PerkinElmer) by exciting Cy5 at 640 nm and measuring emission at 680 nm. Fluorescence signal intensity was quantified using the Living Image software (PerkinElmer).

### Statistical analysis

PCA was performed in the R statistical environment (https://www.r-project.org/) using the prcomp function. Hierarchical clustering was performed using GENE-E (Broad Institute; https://software.broadinstitute.org/GENE-E/). All remaining statistical analyses were conducted in GraphPad Prism 8.0. All sample sizes, statistical tests, and *p*-values are specified in figure legends.

## AUTHOR CONTRIBUTIONS

Conceptualization, A.P.S., J.D.K., J.S.D., and S.N.B.; Methodology, A.P.S. and J.D.K.; Data Curation, Validation, and Formal Analysis, A.P.S. and J.D.K.; Investigation, A.P.S., J.D.K., S.S., Q.Z., and A.B.; Visualization, A.P.S. and J.D.K.; Writing – Original Draft, A.P.S., J.D.K., and S.N.B.; Writing – Review & Editing, A.P.S., J.D.K., S.S., J.S.D., Q.Z., A.B., and S.N.B.; Resources and Funding Acquisition, S.N.B.; Project Administration and Supervision, A.P.S., J.D.K., and S.N.B.; A.P.S. and J.D.K. contributed equally.

## ACKNOWLEDGEMENTS

We thank Dr. H. Fleming (MIT) for critical reading and editing of the manuscript; Dr. J. Zhao, Dr. P. Watson, and Dr. C. Sawyers (Memorial Sloan-Kettering) for valuable technical discussion and insight; M. Anahtar, C. Martin-Alonso, and R. Zhao (MIT) for technical assistance; Dr. P. Blainey (MIT and Broad Institute) and Dr. M. Vander Heiden (MIT) for helpful discussion; and K. Cormier and Dr. R. Bronson of the Koch Institute Histology Core (MIT). This study was supported in part by a Koch Institute Support Grant P30-CA14051 from the National Cancer Institute (Swanson Biotechnology Center), a Core Center Grant P30-ES002109 from the National Institute of Environmental Health Sciences, the Ludwig Fund for Cancer Research, Janssen Pharmaceuticals, and the Koch Institute Marble Center for Cancer Nanomedicine. A.P.S. acknowledges support from the NIH Molecular Biophysics Training Grant (NIH/NIGMS T32 GM008313) and the National Science Foundation Graduate Research Fellowship. J.D.K. acknowledges support from the Ludwig Center fellowship. J.S.D acknowledges support from the National Science Foundation Graduate Research Fellowship, the Ludwig Center fellowship, and the Siebel Scholar Foundation. A.B. acknowledges support from the Early Postdoc Fellowship program (P2ELP2_178238) from Swiss National Science Foundation. S.N.B is a Howard Hughes Medical Institute Investigator.

## CONFLICT OF INTEREST STATEMENT

S.N.B. holds equity in Glympse Bio and Impilo Therapeutics; is a director at Vertex; consults for Cristal, Maverick, and Moderna; and receives sponsored research funds from Johnson & Johnson.

## DATA AVAILABILITY

The data that support the findings of this study are available upon reasonable request to.

## SUPPLEMENTARY FIGURES

**Supplementary Figure S1.**
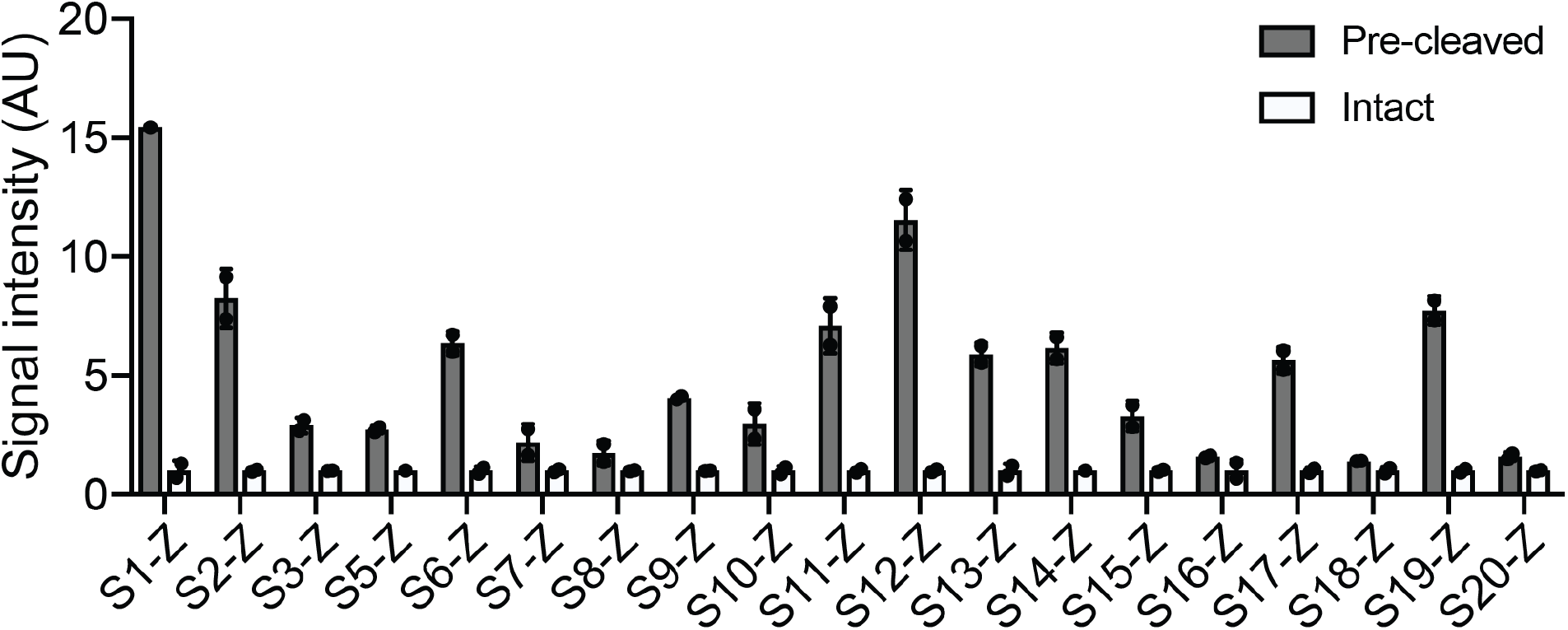
AZP library characterization. AZPs, either intact or with linkers pre-cleaved by a cognate recombinant protease, were applied to fresh frozen colon tissue for 30 minutes, and fluorescent signal intensity of bound probes was quantified (*n* = 1-2 replicates per probe; mean ± s.d.).

**Supplementary Figure S2.**
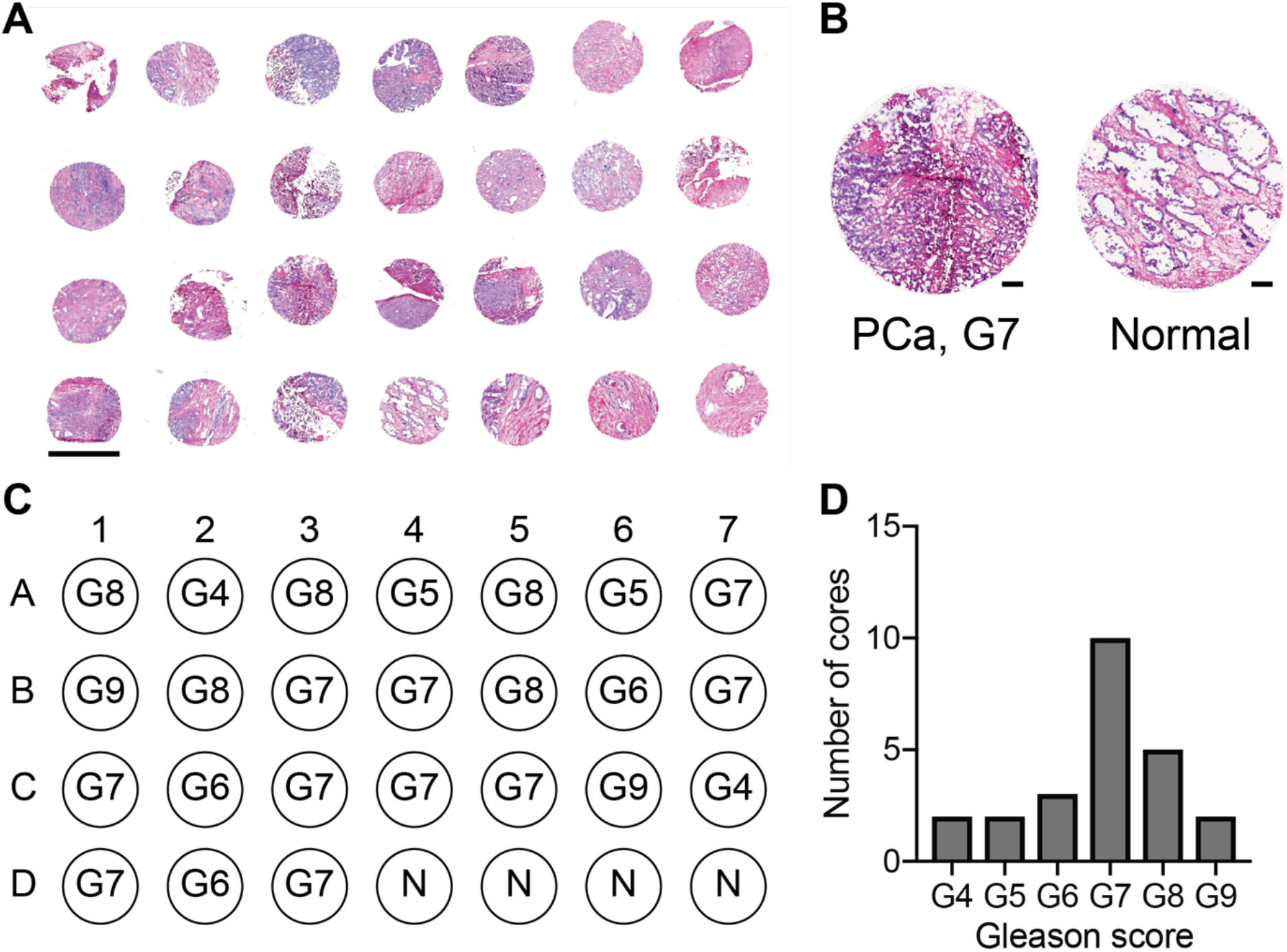
Fresh frozen human prostate cancer tissue microarray (TMA). (a) H&E stain of human prostate cancer (PCa) TMA. Scale bar = 2 mm. (b) Hematoxylin and eosin stain of select biopsy cores from Gleason 7 PCa tumor (left) and normal prostate (right). Scale bars = 200 μm. (c) TMA map detailing the Gleason scores (i.e., G4-G10) for prostate cancer specimens. N = normal. (d) Distribution across Gleason scores for cores in the TMA.

**Supplementary Figure S3.**
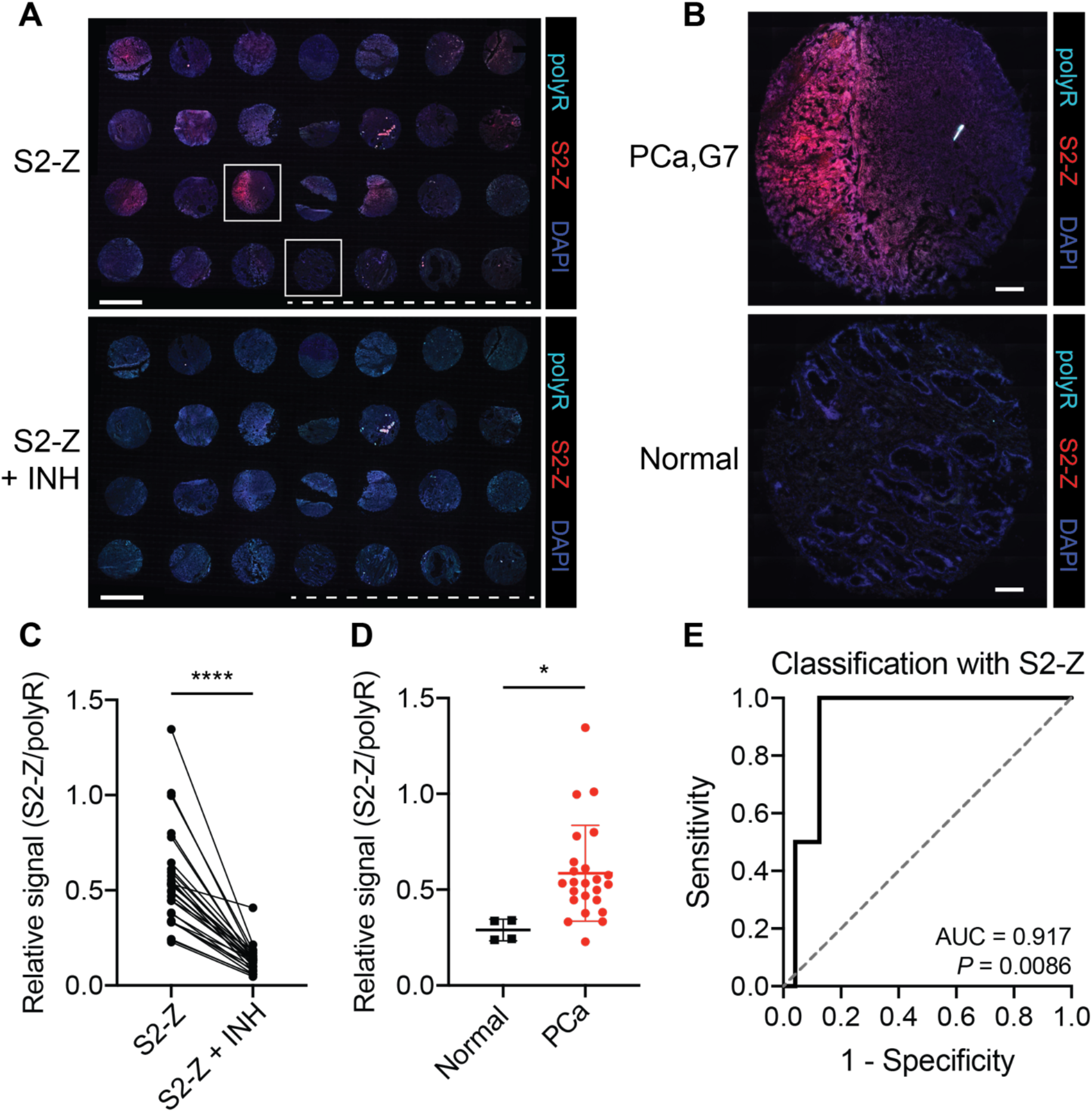
S2-Z selectively labels human PCa tissue. (a) Application of S2-Z AZP (red) to a human PCa tissue microarray (TMA) consisting of 24 prostate adenocarcinoma samples and 4 normal prostate samples (S2-Z, top). A consecutive TMA was stained with S2-Z along with a cocktail of protease inhibitors (S2-Z + INH, bottom). Sections were stained with a polyR binding control (teal) and counterstained with DAPI (blue). Dotted lines are shown below normal prostate samples. Scale bars = 2 mm. (b) Higher-magnification image of boxed cores from showing Gleason 7 PCa (top) and normal prostate (bottom). Scale bars = 200 μm. (c) Quantification of average S2-Z intensity relative to polyR (binding control) intensity across each TMA core (*n* = 28) for sections incubated with (S2-Z + INH) and without (S2-Z) protease inhibitors (two-tailed paired *t*-test, \*\*\*\**P* < 0.0001). (d) Quantification of relative S2-Z intensity from normal (*n* = 4) and PCa tumor (*n* = 24) cores (mean ± s.d.; two-tailed unpaired *t*-test, \**P* = 0.0284). (e) Receiver-operating characteristic (ROC) curve showing performance of relative AZP signal (S2-Z/polyR) in discriminating normal from PCa tumor cores (AUC = 0.917, 95% confidence interval 0.8103-1.000; *P* = 0.0086 from random classifier shown in dashed line).

**Supplementary Figure S4.**
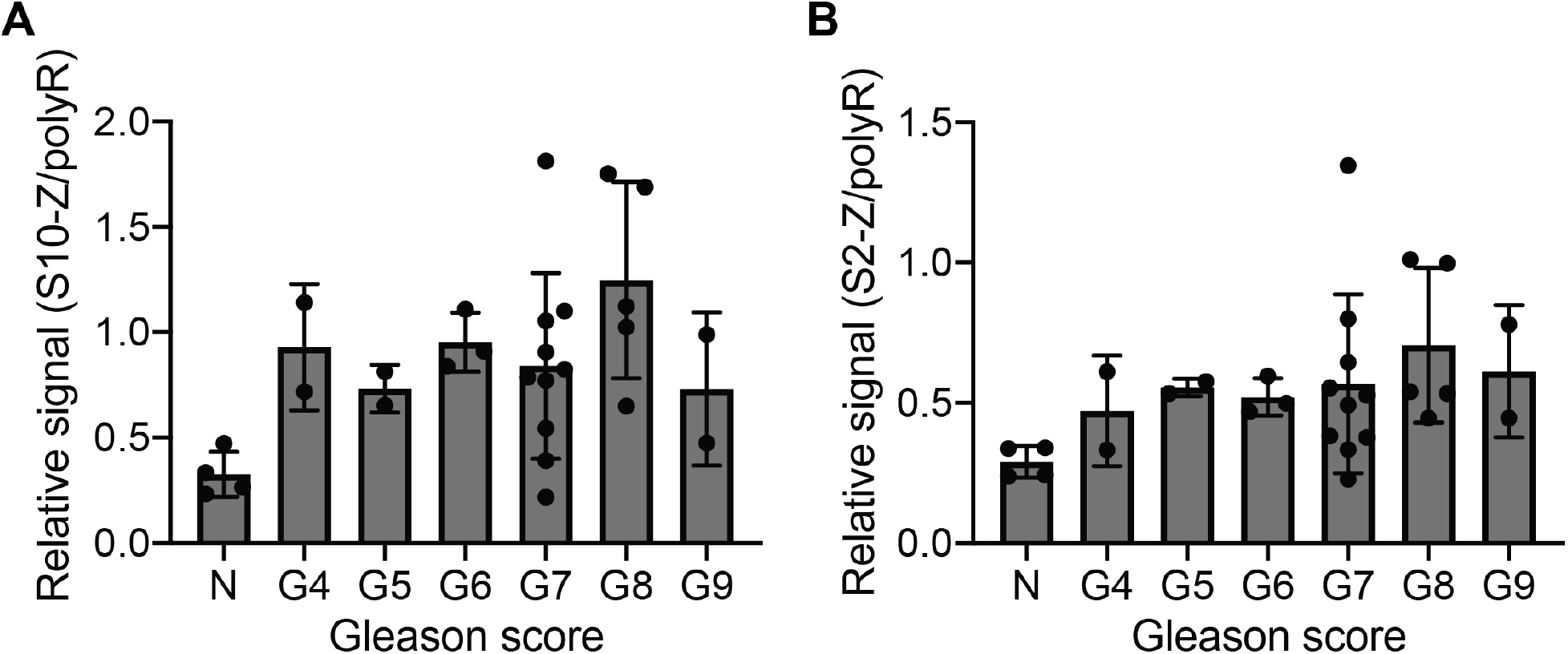
Distribution of AZP staining intensities in human PCa TMA. Distribution of relative S10-Z (a) and S2-Z (b) intensities across Gleason scores across the TMA (N = normal; AZP intensities normalized to polyR binding control).

**Supplementary Figure S5.**
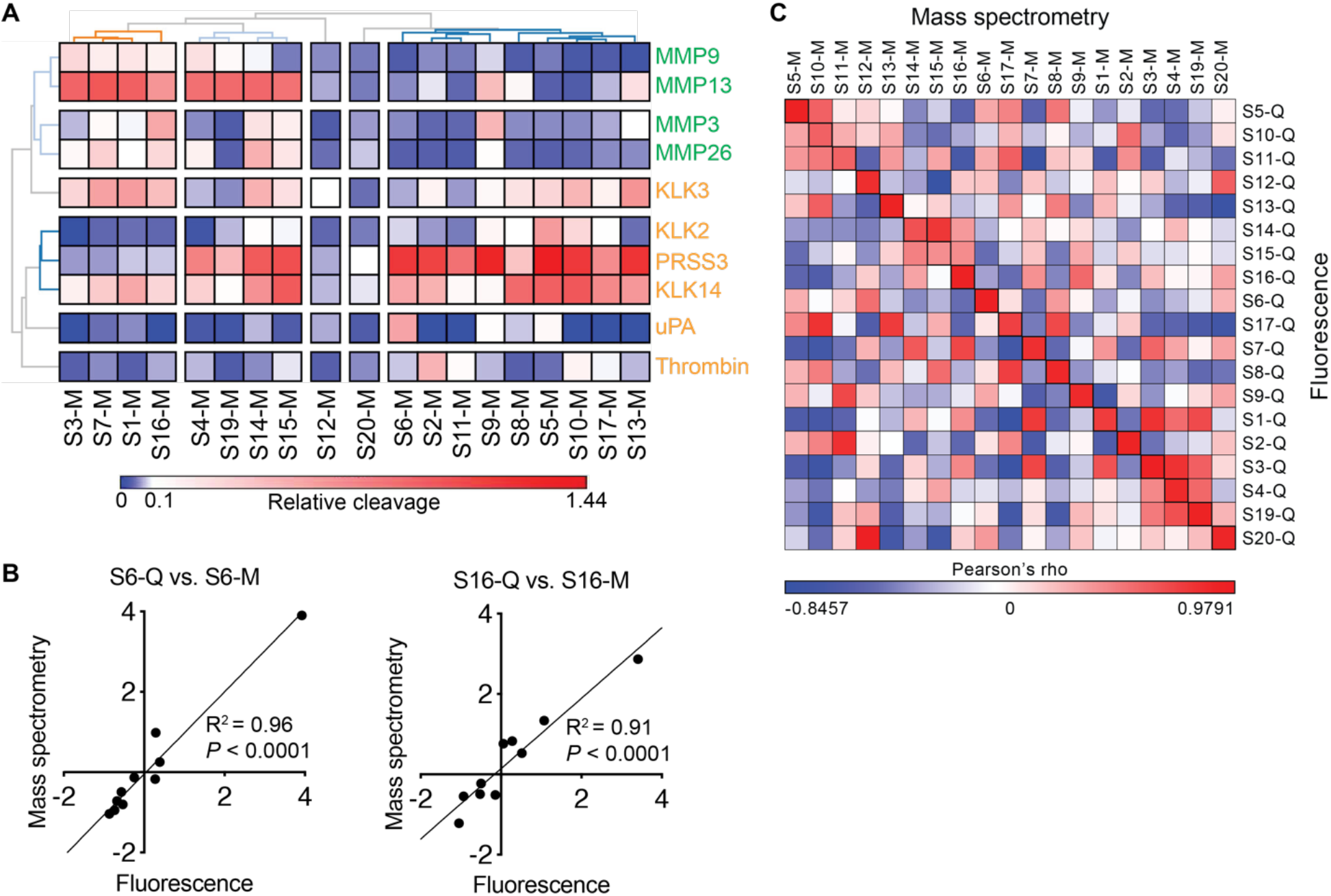
Protease activity profiling with barcoded substrate libraries. (a) Heat map showing *in vitro* cleavage of mass-barcoded substrates (x-axis) by selected recombinant proteases (y-axis; metalloproteinases, green; serine proteases, yellow). Cleavage products were quantified by mass spectrometry, and unsupervised hierarchical clustering was performed (*n* = 2 replicates per protease). (b) Comparison of relative substrate cleavage *z*-scores for serine protease substrate S6-Q/S6-M (quenched/mass encoded, respectively) and metalloproteinase substrate S16-Q/S16-M (quenched/mass encoded, respectively), as measured by fluorescence with quenched substrates (x-axis) and mass spectrometry with bead-conjugated substrates (y-axis) (linear regression; S6 R^2^ = 0.96, *P* < 0.0001 from non-zero slope; S16 R^2^ = 0.91, *P* < 0.0001 from non-zero slope). (c) Correlation of cleavage *z-*scores of FRET-paired free peptides to cleavage z-scores of mass-barcoded peptides conjugated to the surface of magnetic beads, calculated as Pearson’s rho across all proteases using cleavage *z*-scores.

**Supplementary Figure S6.**
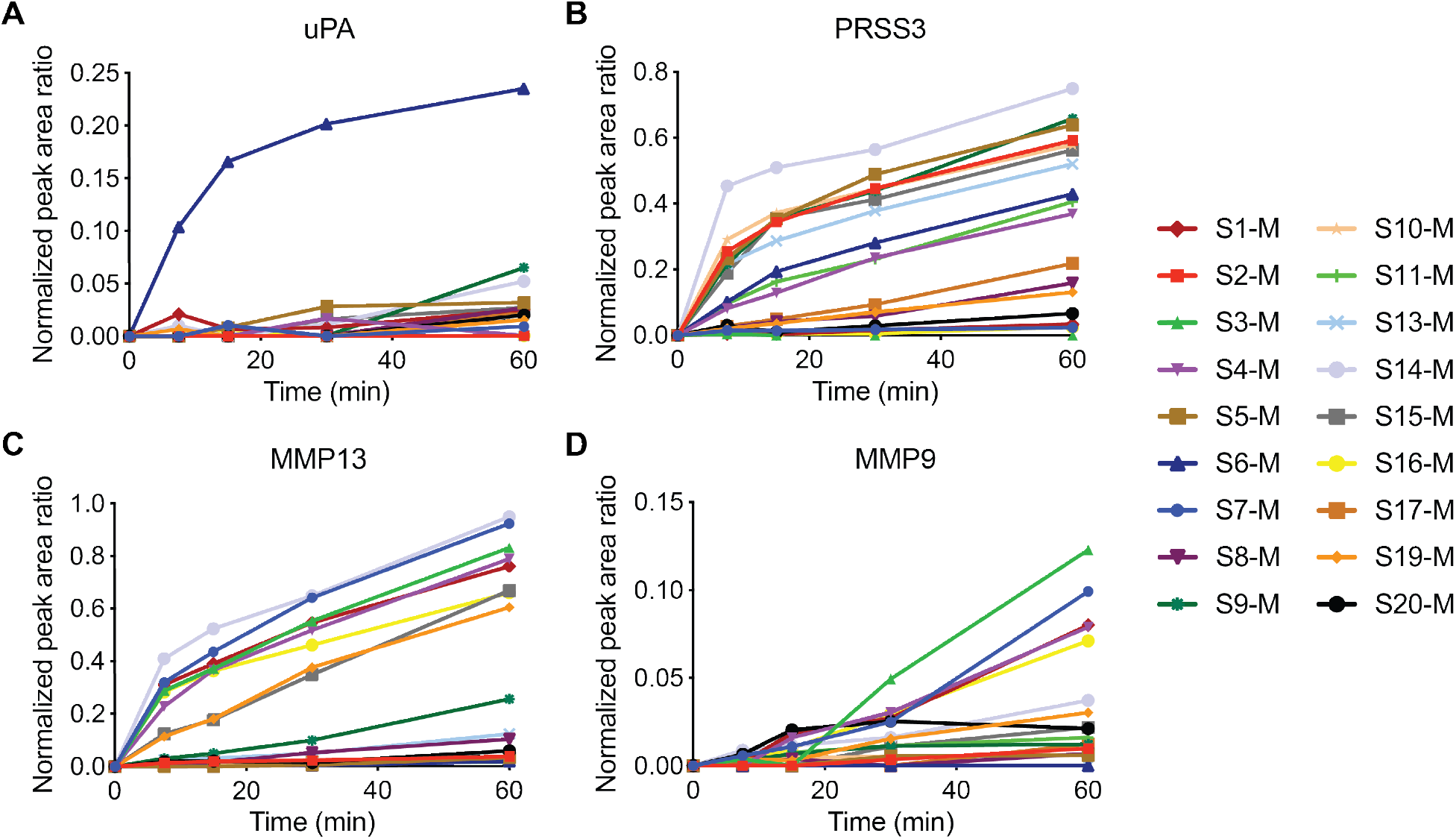
Profiling cleavage kinetics using mass-barcoded bead library. Kinetics of substrate cleavage obtained from mass-barcoded library screen for the serine proteases uPA (a) and PRSS3 (b) and the metalloproteinases MMP13 (c) and MMP9 (d). Kinetics were assessed via quantification of liberated barcodes isolated via magnetic separation at various time points after addition of protease, followed by mass spectrometry quantification with LC-MS/MS. Lines represent means of two independent measurements.

**Supplementary Figure S7.**
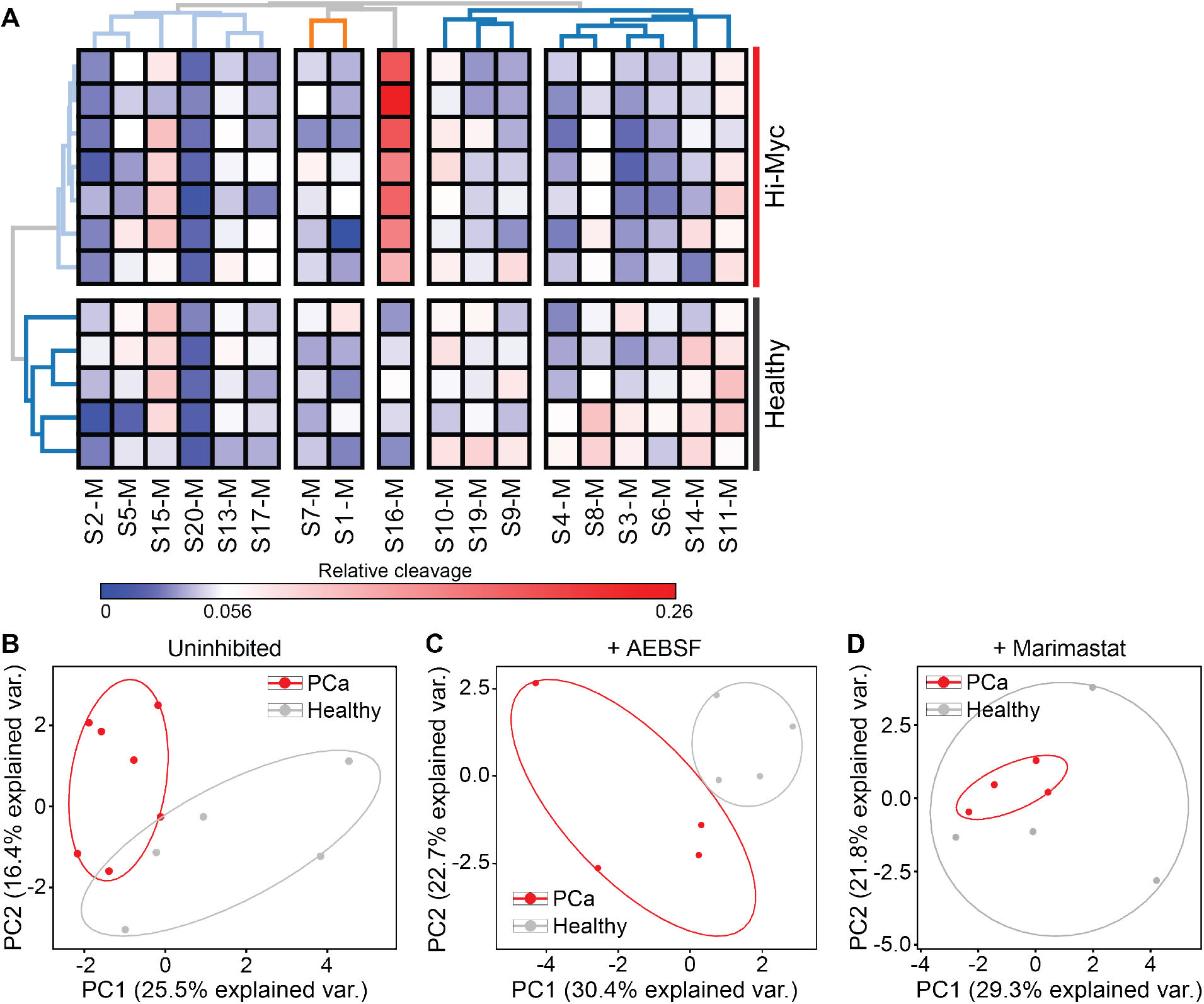
Differential cleavage of peptide S16 is driven by MMP dysregulation and drives differentiation of Hi-Myc from healthy prostates. (a) Hierarchical clustering of cleavage data from multiplexed protease substrate screen of mass-encoded bead library against homogenates of prostates from healthy (gray, *n* = 5) and Hi-Myc (red, *n* = 7) mice. (b-d) Principal component analysis (PCA) of cleavage data from homogenates incubated without inhibitor (b), with the serine protease inhibitor AEBSF (c), or with the metalloproteinase inhibitor marimastat (d).

**Supplementary Figure S8.**
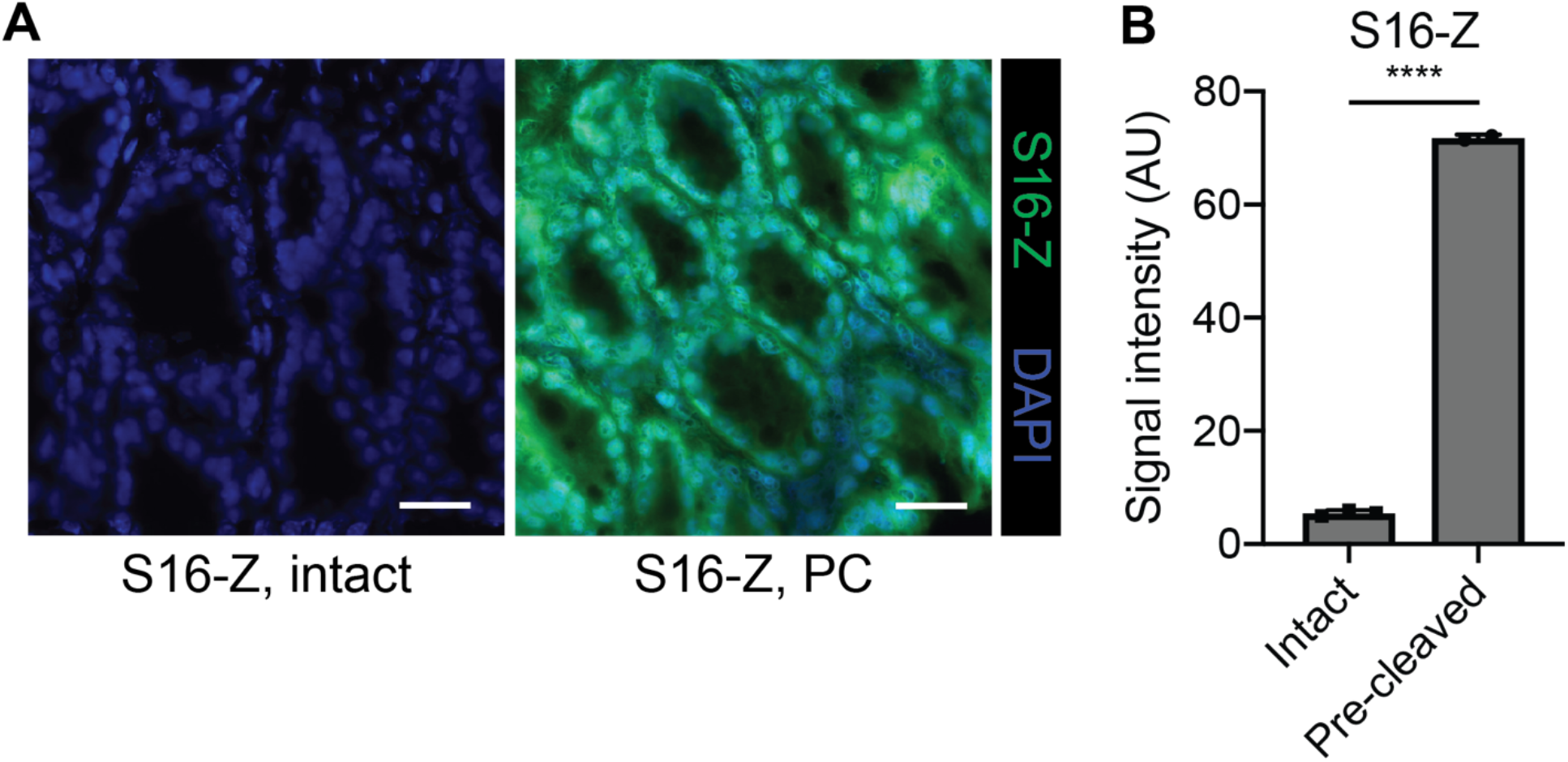
S16-Z tissue binding depends on proteolytic cleavage. (a) Binding of intact or MMP12 pre-cleaved (PC) S16-Z (green) to fresh frozen mouse colon tissue following incubation at 4° C. Sections were counterstained with DAPI (blue). Scale bars = 25 μm. Quantification of S16-Z binding for intact probe or probe pre-cleaved by MMP12 (*n* = 2-3 replicates; mean ± s.d.; two-tailed unpaired *t*-test, \*\*\*\**P* < 0.0001).

**Supplementary Figure S9.**
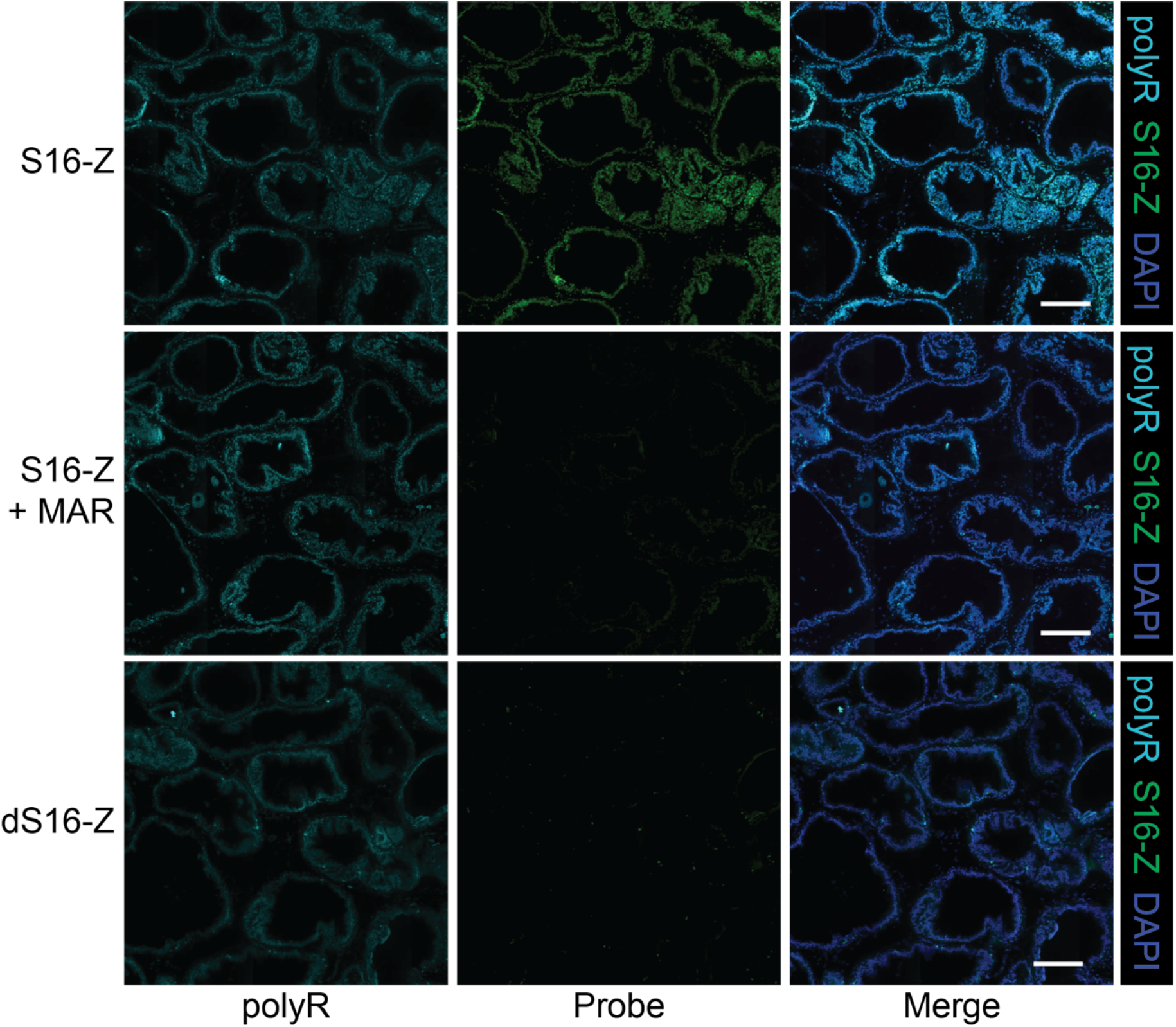
S16-Z labeling of Hi-Myc tissue is dependent on *in situ* MMP activity. Staining of Hi-Myc tissue with polyR (left column, teal) and either MMP-activatable S16-Z (top and middle rows; green) or *d*-stereoisomer dS16-Z (bottom row; green). Top and middle rows show staining of consecutive sections without (top) and with (middle) the MMP inhibitor marimastat (MAR). Sections were counterstained with DAPI (blue). Scale bars = 200 μm.

**Supplementary Figure S10.**
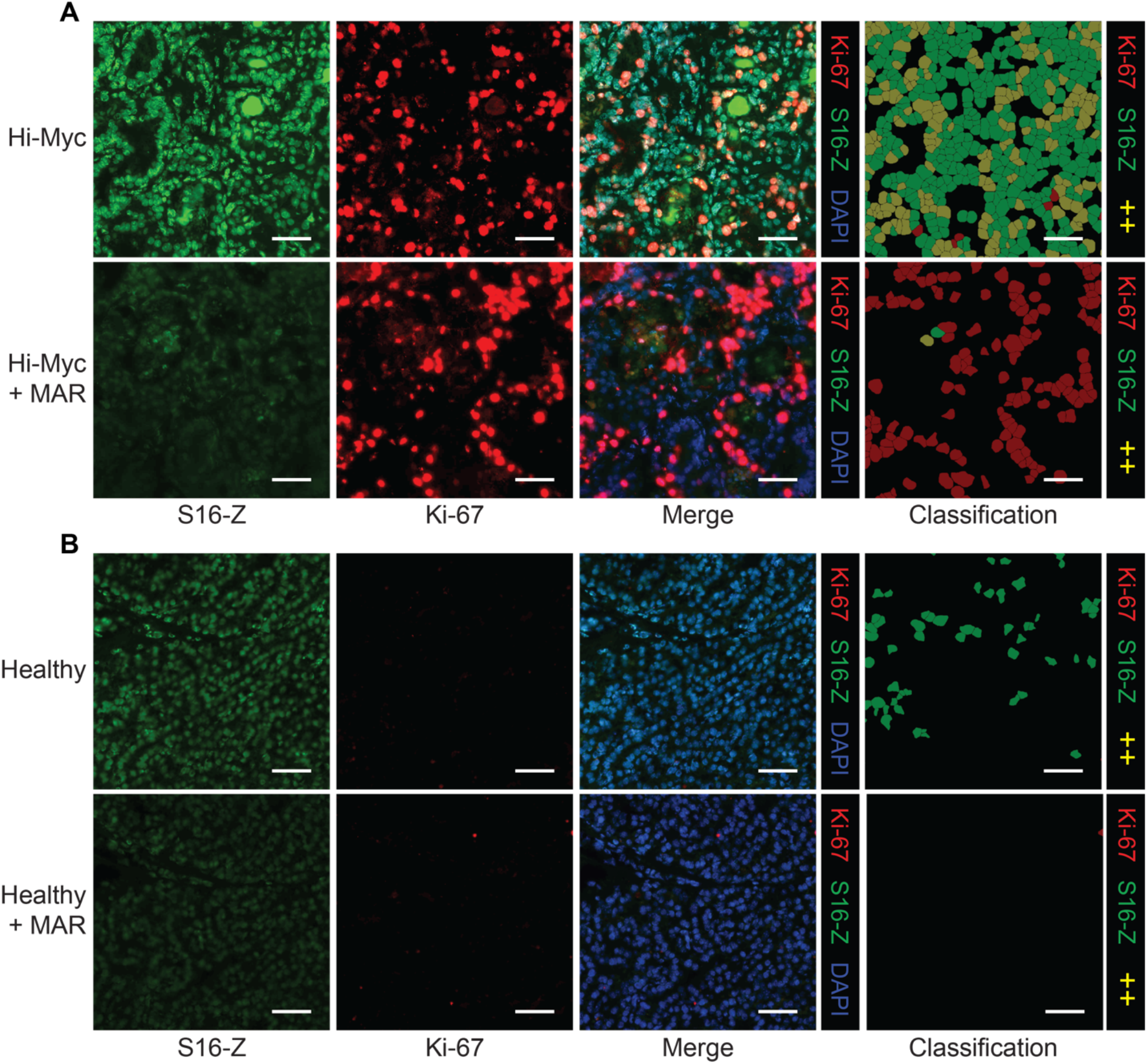
MMP activity drives S16-Z labeling of proliferative tumor regions in Hi-Myc prostates. (a, b) Staining of Hi-Myc tumor region (a) and healthy prostate tissue (b) with S16-Z (green) with co-staining for the proliferation marker Ki-67 (red). Consecutive sections were stained with S16-Z and Ki-67 in the presence of marimastat (MAR). Sections were counterstained with DAPI (blue). Detected cells were classified on the basis of S16-Z and Ki-67 staining intensities to produce cellular classification maps (green: S16-Z+, red: Ki67+, yellow: S16-Z+ and Ki-67+). Scale bars = 50 μm.

**Supplementary Figure S11.**
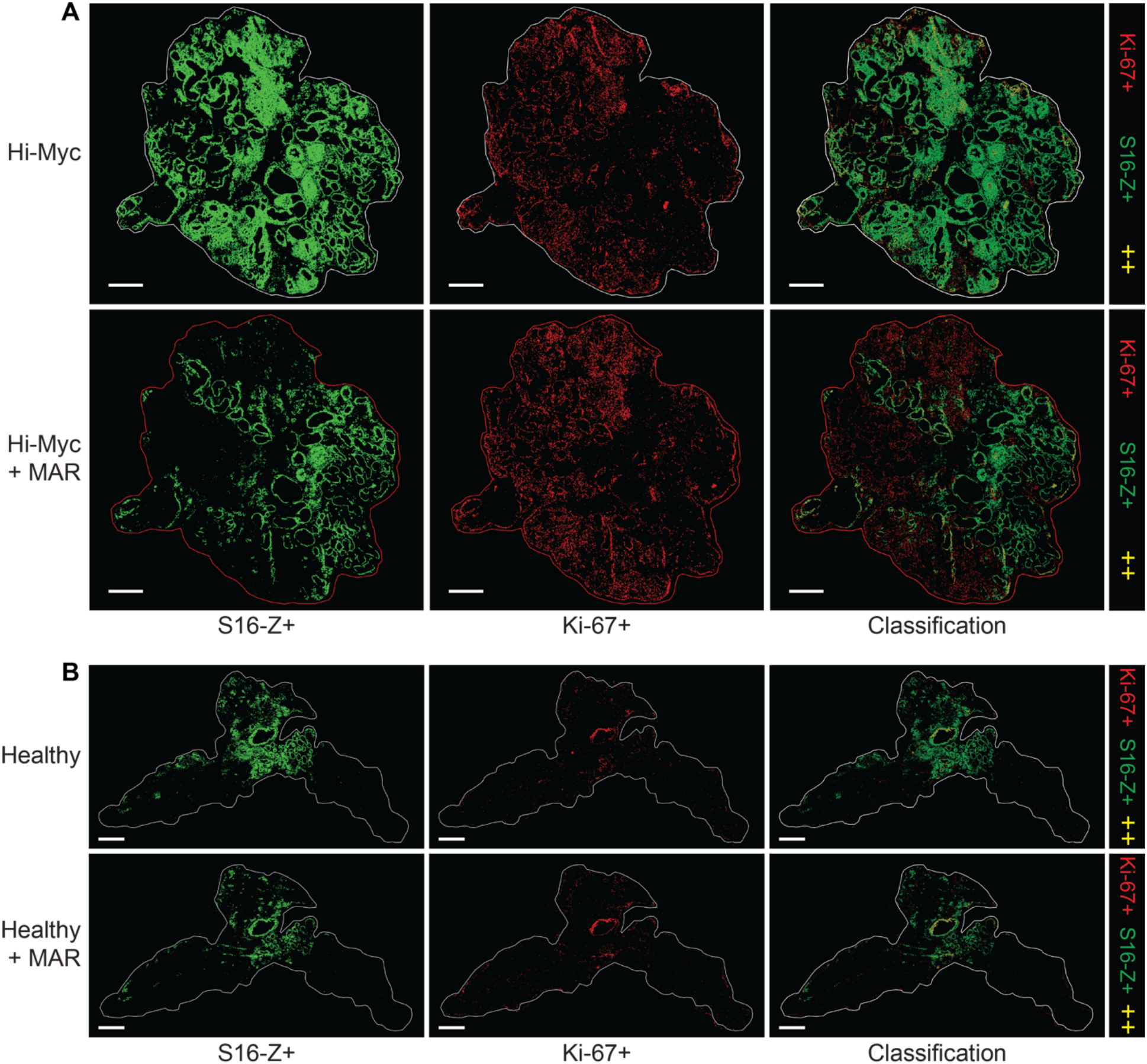
S16-Z and Ki-67 staining of healthy and Hi-Myc prostate whole tissue sections. (a, b) S16-Z, with or without MAR, was applied to prostate tissues from Hi-Myc and healthy mice with co-staining for the proliferation marker Ki-67. Detected cells were classified on the basis of S16-Z and Ki-67 staining intensities to produce classification maps of whole tissue sections (green: S16-Z+, red: Ki67+, yellow: S16-Z+ and Ki-67+). Scale bars = 1 mm.

**Supplementary Figure S12.**
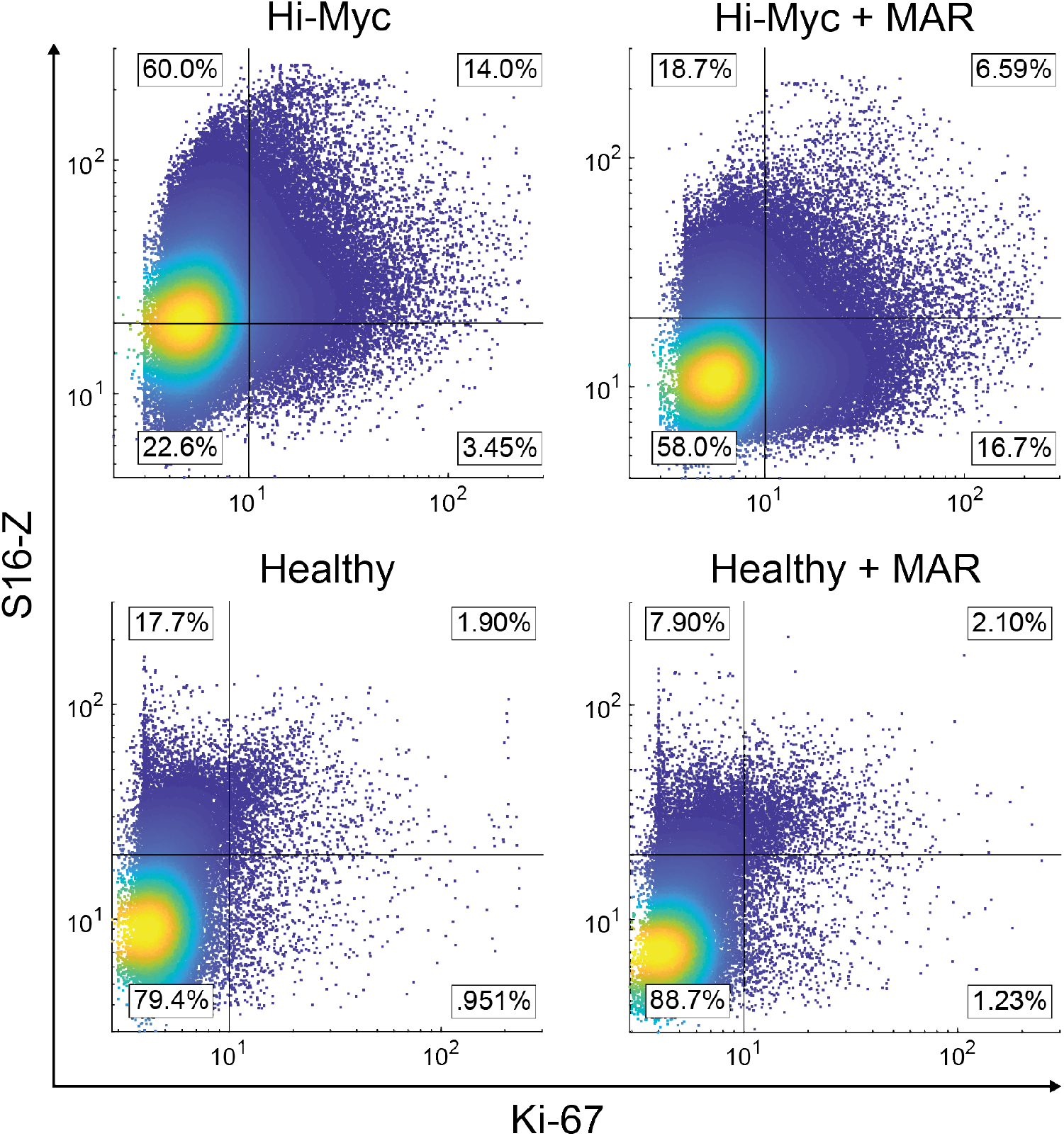
Whole sample, cell-by-cell quantification of S16-Z and Ki-67 fluorescent staining. Cell-by-cell quantification of S16-Z and Ki-67 fluorescence intensity in detected nuclei from representative Hi-Myc and healthy prostate tissue sections incubated with or without the MMP inhibitor marimastat (MAR). Lines represent thresholds for positivity for each of the markers.

**Supplementary Figure S13.**
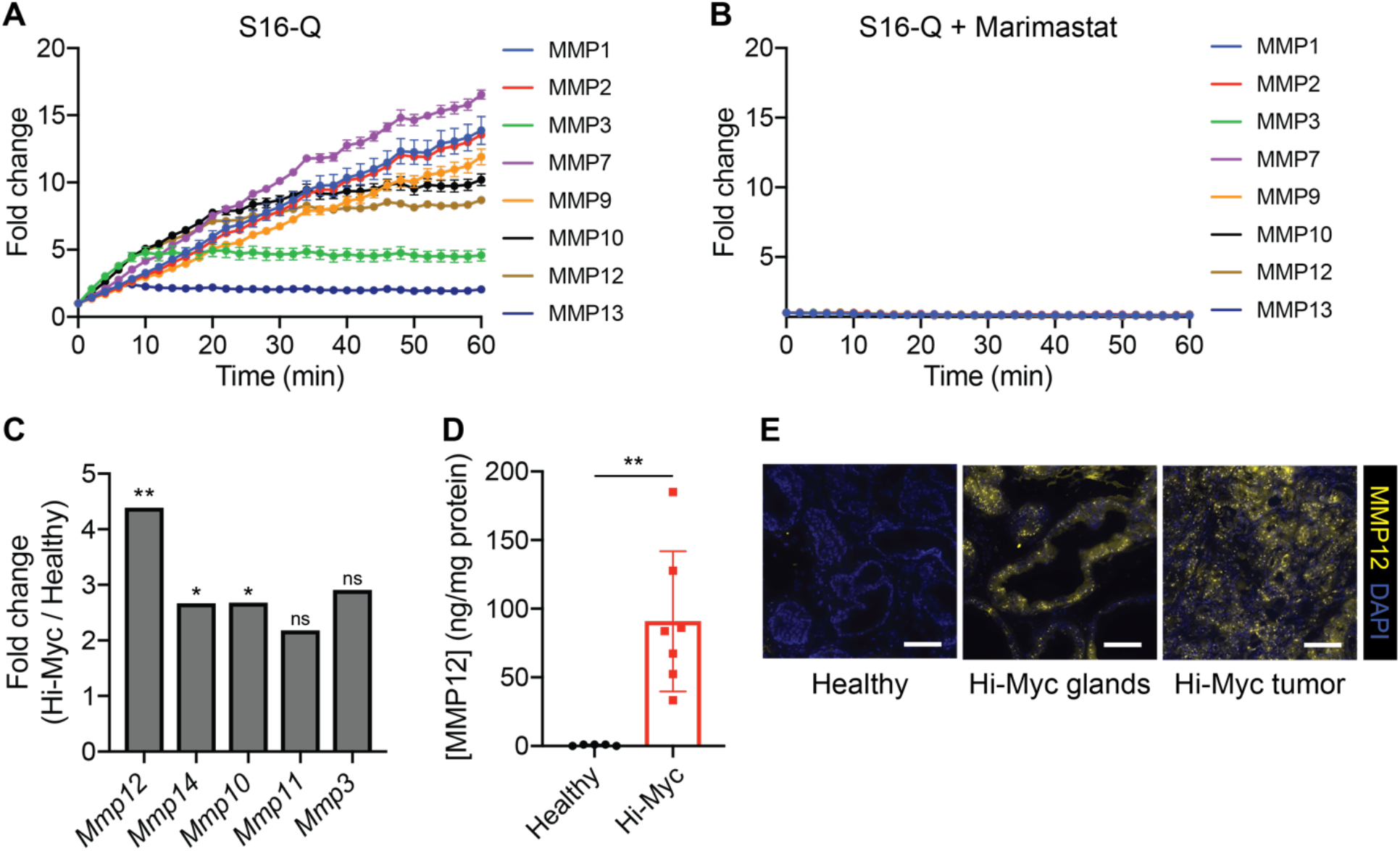
Characterization of MMP12 dysregulation in Hi-Myc model. (a) Fluorescent dequenching measurements for panel of MMPs against fluorescent version (S16-Q) of Hi-Myc-responsive substrate S16 (*n* = 2 replicates per protease; mean ± s.d.). (b) Addition of MMP inhibitor marimastat (MAR) abrogated fluorescence increase for panel of MMPs against S16-Q (*n* = 2 replicates per protease; mean ± s.d.). (c) Relative expression of the top 5 MMPs most significantly upregulated in Hi-Myc prostates relative to age-matched healthy controls, analyzed by Affymetrix microarray, adapted from Ellwood-Yen et al., *Cancer Cell* 2003 (35) (two-tailed unpaired *t*-test; \*\**P* < 0.01, \**P* < 0.05, ns = not significant). (d) Enzyme-linked immunosorbent assay (ELISA) results comparing protein-level abundance of MMP12 in homogenates of Hi-Myc prostates relative to age-matched healthy controls (*n* = 7 Hi-Myc, *n* = 5 healthy; mean ± s.d.; Welch’s unequal variances *t*-test; \*\**P* = 0.0034). (e) Immunofluorescence staining for MMP12 (yellow) in fresh-frozen sections of healthy and Hi-Myc prostates, showing histologically similar regions of healthy and Hi-Myc prostates (Healthy, Hi-Myc glands) as well as a Hi-Myc tumor region (Hi-Myc tumor). Scale bars = 100 μm.

**Supplementary Table S1.**
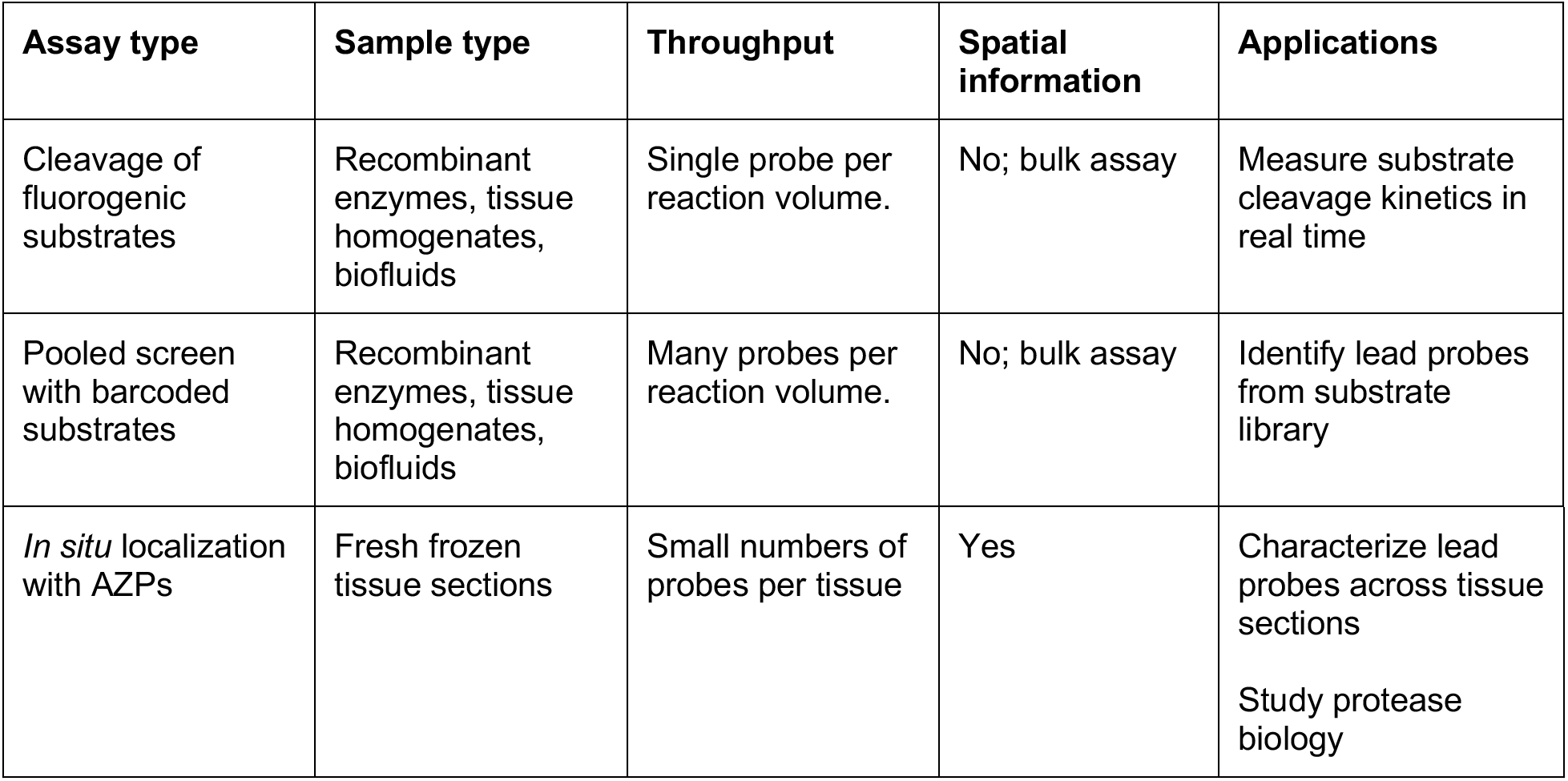
Strategies for profiling protease activity *ex vivo*.

## REFERENCES

1. Dudani JS, Warren AD, Bhatia SN. Harnessing Protease Activity to Improve Cancer Care. Annu Rev Cancer Biol. 2018;2:353–76.

2. Kessenbrock K, Plaks V, Werb Z. Matrix Metalloproteinases: Regulators of the Tumor Microenvironment. Cell. 2010;141:52–67.

3. Soleimany AP, Bhatia SN. Activity-Based Diagnostics: An Emerging Paradigm for Disease Detection and Monitoring. Trends Mol Med. 2020;26:450–68.

4. Weissleder R, Tung C-H, Mahmood U, Bogdanov A. In vivo imaging of tumors with protease-activated near-infrared fluorescent probes. Nat Biotechnol. 1999;17:375–8.

5. Sanman LE, Bogyo M. Activity-Based Profiling of Proteases. Annu Rev Biochem. 2014;83:249–73.

6. Blum G, Von Degenfeld G, Merchant MJ, Blau HM, Bogyo M. Noninvasive optical imaging of cysteine protease activity using fluorescently quenched activity-based probes. Nat Chem Biol. 2007;3:668–77.

7. Kwong GA, Von Maltzahn G, Murugappan G, Abudayyeh O, Mo S, Papayannopoulos IA, et al. Mass-encoded synthetic biomarkers for multiplexed urinary monitoring of disease. Nat Biotechnol. 2013;31:63–70.

8. Kwon EJ, Dudani JS, Bhatia SN. Ultrasensitive tumour-penetrating nanosensors of protease activity. Nat Biomed Eng. 2017;1:1–10.

9. Loynachan CN, Soleimany AP, Dudani JS, Lin Y, Najer A, Bekdemir A, et al. Renal clearable catalytic gold nanoclusters for in vivo disease monitoring. Nat Nanotechnol. Springer US; 2019;14:883–90.

10. Kirkpatrick JD, Warren AD, Soleimany AP, Westcott PMK, Voog JC, Martin-alonso C, et al. Urinary detection of lung cancer in mice via noninvasive pulmonary protease profiling. Sci Transl Med. 2020;12:eaaw0262.

11. Desnoyers LR, Vasiljeva O, Richardson JH, Yang A, Menendez EEM, Liang TW, et al. Tumor-specific activation of an EGFR-targeting probody enhances therapeutic index. Sci Transl Med. 2013;5:207ra144.

12. Polu KR, Lowman HB. Probody therapeutics for targeting antibodies to diseased tissue. Expert Opin Biol Ther. 2014;14:1049–53.

13. Chen IJ, Chuang CH, Hsieh YC, Lu YC, Lin WW, Huang CC, et al. Selective antibody activation through protease-activated pro-antibodies that mask binding sites with inhibitory domains. Sci Rep. 2017;7:1–12.

14. Trang VH, Zhang X, Yumul RC, Zeng W, Stone IJ, Wo SW, et al. A coiled-coil masking domain for selective activation of therapeutic antibodies. Nat Biotechnol. 2019;37:761–5.

15. Wang D, Wang T, Yu H, Feng B, Zhou L, Zhou F, et al. Engineering nanoparticles to locally activate T cells in the tumor microenvironment. Sci Immunol. 2019;4:eaau6584.

16. Millar DG, Ramjiawan RR, Kawaguchi K, Gupta N, Chen J, Zhang S, et al. Antibody-mediated delivery of viral epitopes to tumors harnesses CMV-specific T cells for cancer therapy. Nat Biotechnol. 2020;38:420–5.

17. Withana NP, Ma X, Mcguire HM, Verdoes M, Van Der Linden WA, Ofori LO, et al. Non-invasive Imaging of Idiopathic Pulmonary Fibrosis Using Cathepsin Protease Probes. Sci Rep. 2015;6:19755.

18. Nguyen QT, Olson ES, Aguilera TA, Jiang T, Scadeng M, Ellies LG, et al. Surgery with molecular fluorescence imaging using activatable cell-penetrating peptides decreases residual cancer and improves survival. Proc Natl Acad Sci U S A. 2010;107:4317–22.

19. Olson ES, Jiang T, Aguilera TA, Nguyen QT, Ellies LG, Scadeng M, et al. Activatable cell penetrating peptides linked to nanoparticles as dual probes for in vivo fluorescence and MR imaging of proteases. Proc Natl Acad Sci U S A. 2010;107:4311–6.

20. Whitley MJ, Cardona DM, Lazarides AL, Spasojevic I, Ferrer JM, Cahill J, et al. A mouse-human phase 1 co-clinical trial of a protease-activated fluorescent probe for imaging cancer. Sci Transl Med. 2016;8:320ra4.

21. Mahrus S, Trinidad JC, Barkan DT, Sali A, Burlingame AL, Wells JA. Global sequencing of proteolytic cleavage sites in apoptosis by specific labeling of protein N termini. Cell. 2008;134:866–76.

22. Kleifeld O, Doucet A, Auf Dem Keller U, Prudova A, Schilling O, Kainthan RK, et al. Isotopic labeling of terminal amines in complex samples identifies protein N-termini and protease cleavage products. Nat Biotechnol. 2010;28:281–8.

23. Agard NJ, Mahrus S, Trinidad JC, Lynn A, Burlingame AL, Wells JA. Global kinetic analysis of proteolysis via quantitative targeted proteomics. Proc Natl Acad Sci U S A. 2012;109:1913–8.

24. Villanueva J, Shaffer DR, Philip J, Chaparro CA, Erdjument-Bromage H, Olshen AB, et al. Differential exoprotease activities confer tumor-specific serum peptidome patterns. J Clin Invest. 2006;116:271–84.

25. Chen CH, Miller MA, Sarkar A, Beste MT, Isaacson KB, Lauffenburger DA, et al. Multiplexed protease activity assay for low-volume clinical samples using droplet-based microfluidics and its application to endometriosis. J Am Chem Soc. 2013;135:1645–8.

26. O’Donoghue AJ, Eroy-Reveles AA, Knudsen GM, Ingram J, Zhou M, Statnekov JB, et al. Global identification of peptidase specificity by multiplex substrate profiling. Nat Methods. 2012;9:1095–100.

27. Ivry SL, Sharib JM, Dominguez DA, Roy N, Hatcher SE, Yip-Schneider MT, et al. Global protease activity profiling provides differential diagnosis of pancreatic cysts. Clin Cancer Res. 2017;23:4865–74.

28. Vandooren J, Geurts N, Martens E, Van Den Steen PE, Opdenakker G. Zymography methods for visualizing hydrolytic enzymes. Nat Methods. 2013;10:211–20.

29. Vasiljeva O, Menendez E, Nguyen M, Craik CS, Michael Kavanaugh W. Monitoring protease activity in biological tissues using antibody prodrugs as sensing probes. Sci Rep. 2020;10:5894.

30. Withana NP, Garland M, Verdoes M, Ofori LO, Segal E, Bogyo M. Labeling of active proteases in fresh-frozen tissues by topical application of quenched activity-based probes. Nat Protoc. 2016;11:184–91.

31. Jiang T, Olson ES, Nguyen QT, Roy M, Jennings PA, Tsien RY. Tumor imaging by means of proteolytic activation of cell-penetrating peptides. Proc Natl Acad Sci U S A. 2004;101:17867–72.

32. Vergnolle N. Protease inhibition as new therapeutic strategy for GI diseases. Gut. 2016;65:1215–24.

33. Uhlén M, Fagerberg L, Hallström BM, Lindskog C, Oksvold P, Mardinoglu A, et al. Tissue-based map of the human proteome. Science. 2015;347:1260419.

34. Dudani JS, Ibrahim M, Kirkpatrick J, Warren AD, Bhatia SN. Classification of prostate cancer using a protease activity nanosensor library. Proc Natl Acad Sci U S A. 2018;115:8954–9.

35. Ellwood-Yen K, Graeber TG, Wongvipat J, Iruela-Arispe ML, Zhang J, Matusik R, et al. Myc-driven murine prostate cancer shares molecular features with human prostate tumors. Cancer Cell. 2003;4:223–38.

36. Scott AM, Wolchok JD, Old LJ. Antibody therapy of cancer. Nat Rev Cancer. 2012;12:278–87.

37. Scholzen T, Gerdes J. The Ki-67 protein: From the known and the unknown. J Cell Physiol. 2000;182:311–22.

38. López-Otín C, Matrisian LM. Emerging roles of proteases in tumor suppression. Nat Rev Cancer. 2007;7:800–8.

39. Dong Z, Kumar R, Yang X, Fidler IJ. Macrophage-derived metalloelastase is responsible for the generation of angiostatin in Lewis lung carcinoma. Cell. 1997;88:801–10.

40. Zhou J, Li S, Leung KK, Zou JY, Derisi JL, Wells A, et al. Deep profiling of protease substrate specificity enabled by dual random and scanned human proteome substrate phage libraries. bioRxiv. 2020;1–73.

41. Kasperkiewicz P, Poreba M, Snipas SJ, Parker H, Winterbourn CC, Salvesen GS, et al. Design of ultrasensitive probes for human neutrophil elastase through hybrid combinatorial substrate library profiling. Proc Natl Acad Sci U S A. 2014;111:2518–23.

42. Poreba M, Salvesen GS, Drag M. Synthesis of a HyCoSuL peptide substrate library to dissect protease substrate specificity. Nat Protoc. 2017;12:2189–214.

43. Bankhead P, Loughrey MB, Fernández JA, Dombrowski Y, McArt DG, Dunne PD, et al. QuPath: Open source software for digital pathology image analysis. Sci Rep. 2017;7:1–7.

